# Machine learning using intrinsic genomic signatures for rapid classification of novel pathogens: COVID-19 case study

**DOI:** 10.1101/2020.02.03.932350

**Authors:** Gurjit S. Randhawa, Maximillian P.M. Soltysiak, Hadi El Roz, Camila P.E. de Souza, Kathleen A. Hill, Lila Kari

## Abstract

As of February 20, 2020, the 2019 novel coronavirus (renamed to COVID-19) spread to 30 countries with 2130 deaths and more than 75500 confirmed cases. COVID-19 is being compared to the infamous SARS coronavirus, which resulted, between November 2002 and July 2003, in 8098 confirmed cases worldwide with a 9.6% death rate and 774 deaths. Though COVID-19 has a death rate of 2.8% as of 20 February, the 75752 confirmed cases in a few weeks (December 8, 2019 to February 20, 2020) are alarming, with cases likely being under-reported given the comparatively longer incubation period. Such outbreaks demand elucidation of taxonomic classification and origin of the virus genomic sequence, for strategic planning, containment, and treatment. This paper identifies an intrinsic COVID-19 genomic signature and uses it together with a machine learning-based alignment-free approach for an ultra-fast, scalable, and highly accurate classification of whole COVID-19 genomes. The proposed method combines supervised machine learning with digital signal processing for genome analyses, augmented by a decision tree approach to the machine learning component, and a Spearman’s rank correlation coefficient analysis for result validation. These tools are used to analyze a large dataset of over 5000 unique viral genomic sequences, totalling 61.8 million bp. Our results support a hypothesis of a bat origin and classify COVID-19 as *Sarbecovirus*, within *Betacoronavirus*. Our method achieves high levels of classification accuracy and discovers the most relevant relationships among over 5,000 viral genomes within a few minutes, *ab initio*, using raw DNA sequence data alone, and without any specialized biological knowledge, training, gene or genome annotations. This suggests that, for novel viral and pathogen genome sequences, this alignment-free whole-genome machine-learning approach can provide a reliable real-time option for taxonomic classification.

## Introduction

Coronaviruses are single-stranded positive-sense RNA viruses that are known to contain some of the largest viral genomes, up to around 32 kbp in length [1–5]. After increases in the number of coronavirus genome sequences available following efforts to investigate the diversity in the wild, the family *Coronaviridae* now contains four genera (International Committee on Taxonomy of Viruses, [6]). While those species that belong to the genera *Alphacoronavirus* and *Betacoronavirus* can infect mammalian hosts, those in *Gammacoronavirus* and the recently defined *Deltacoronavirus* mainly infect avian species [4,7–9]. Phylogenetic studies have revealed a complex evolutionary history, with coronaviruses thought to have ancient origins and recent crossover events that can lead to cross-species infection [8,10–12]. Some of the largest sources of diversity for coronaviruses belong to the strains that infect bats and birds, providing a reservoir in wild animals for recombination and mutation that may enable cross-species transmission into other mammals and humans [4,7,8,10,13].

Like other RNA viruses, coronavirus genomes are known to have genomic plasticity, and this can be attributed to several major factors. RNA-dependent RNA polymerases (RdRp) have high mutation rates, reaching from 1 in 1000 to 1 in 10000 nucleotides during replication [7,14,15]. Coronaviruses are also known to use a template switching mechanism which can contribute to high rates of homologous RNA recombination between their viral genomes [9,16–20]. Furthermore, the large size of coronavirus genomes is thought to be able to accommodate mutations to genes [7]. These factors help contribute to the plasticity and diversity of coronavirus genomes today.

The highly pathogenic human coronaviruses, Severe Acute Respiratory Syndrome coronavirus (SARS-CoV) and Middle East respiratory syndrome coronavirus (MERS-CoV) belong to lineage B (sub-genus *Sarbecovirus)* and lineage C (sub-genus *Merbecovirus)* of *Betacoronavirus*, respectively [9,21–23]. Both result from zoonotic transmission to humans and lead to symptoms of viral pneumonia, including fever, breathing difficulties, and more [24, 25]. Recently, an unidentified pneumonia disease with similar symptoms caused an outbreak in Wuhan and is thought to have started from a local fresh seafood market [26–30]. This was later attributed to a novel coronavirus deemed COVID-19 and represents the third major zoonotic human coronavirus of this century [31]. As of February 20, confirmed cases have risen to 75752 globally, with infections reported in 30 countries [32]. As a result, the World Health Organization set the risk assessment to “Very High” for China, where the bulk of the cases are contained and “High” for regional and global levels [33]. Initiatives to identify the source of transmission and possible intermediate animal vectors have commenced since the genome sequence became publicly available.

From analyses employing whole genome to viral protein-based comparisons, the COVID-19 strain is thought to belong to lineage B *(Sarbecovirus)* of *Betacoronavirus.* From phylogenetic analysis of the RdRp protein, spike proteins, and full genomes of COVID-19 and other coronaviruses, it was found that COVID-19 is most closely related to two bat SARS-like coronaviruses, *bat-SL-CoVZXC21* and *bat-SL-CoVZC45*, found in Chinese horseshoe bats *Rhinolophus sinicus* [12,34–38]. Along with the phylogenetic data, the genome organization of COVID-19 was found to be typical of lineage B *(Sarbecovirus) Betacoronaviruses* [34]. From phylogenetic analysis of full genome alignment and similarity plots, it was found that COVID-19 has the highest similarity to the bat coronavirus *RaTG13* [39]. Close associations to bat coronavirus *RaTG13* and two bat SARS-like CoVs (*ZC45* and *ZXC21*) are also supported in alignment-based phylogenetic analyses [39]. Within the COVID-19 strains, over 99% sequence similarity and a lack of diversity within these strains suggest a common lineage and source, with support for recent emergence of the human strain [12, 31]. There is still ongoing debate whether the COVID-19 strain arose following recombination with previously identified bat and unknown coronaviruses [40] or arose independently as a new lineage to infect humans [39]. In combination with the identification that the angiotensin converting enzyme 2 (ACE2) protein is a receptor for COVID-19, as it is for SARS and other *Sarbecovirus* strains, the hypothesis that COVID-19 originated from bats is deemed very likely [12,34,36,39,42–45].

All analyses performed thus far have been alignment-based and rely on the annotations of the viral genes. Though alignment-based methods have been successful in finding sequence similarities, their application can be challenging in many cases [46, 47]. It is realistically impossible to analyze thousands of complete genomes using alignment-based methods due to the heavy computation time. Moreover, the alignment demands the sequences to be continuously homologous which is not always the case. Alignment-free methods [48–52] have been proposed in the past as an alternative to address the limitations of the alignment-based methods. Comparative genomics beyond alignment-based approaches have benefited from the computational power of machine learning. Machine learning-based alignment-free methods have also been used successfully for a variety of problems including virus classification [50–52]. An alignment-free approach [50] was proposed for subtype classification of HIV-1 genomes and achieved ~ 97% classification accuracy. MLDSP [51], with the use of a broad range of *1D* numerical representations of DNA sequences, has also achieved very high levels of classification accuracy with viruses. Even rapidly evolving, plastic genomes of viruses such as *Influenza* and *Dengue* are classified down to the level of strain and subtype, respectively with 100% classification accuracy. MLDSP-GUI [52] provides an option to use 2D Chaos Game Representation (CGR) [53] as numerical representation of DNA sequences. CGR’s have a longstanding use in species classification with identification of biases in sequence composition [49,52,53]. MLDSP-GUI has shown 100% classification accuracy for *Flavivirus* genus to species classification using *2D* CGR as numerical representation [52]. MLDSP and MLDSP-GUI have demonstrated the ability to identify the genomic signatures (a species-specific pattern known to be pervasive throughout the genome) with species level accuracy that can be used for sequence (dis)similarity analyses. In this study, we use MLDSP [51] and MLDSP-GUI [52] with CGR as a numerical representation of DNA sequences to assess the classification of COVID-19 from the perspective of machine learning-based alignment-free whole genome comparison of genomic signatures. Using MLDSP and MLDSP-GUI, we confirm that the COVID-19 belongs to the *Betacoronavirus*, while its genomic similarity to the sub-genus *Sarbecovirus* supports a possible bat origin.

This paper shows how machine learning using intrinsic genomic signatures can provide rapid alignment-free taxonomic classification of novel pathogens. Our method delivers accurate classifications of COVID-19 without *a priori* biological knowledge, by a simultaneous processing of the geometric space of all relevant viral genomes. The main contributions are:

- Identifying intrinsic viral genomic signatures, and utilizing them for a real-time and highly accurate machine learning-based classification of novel pathogen sequences, such as COVID-19;
- A general-purpose bare-bones approach, which uses raw DNA sequences alone and does not have any requirements for gene or genome annotation;
- The use of a “decision tree” approach to supervised machine learning (paralleling taxonomic ranks), for successive refinements of taxonomic classification.
- A comprehensive and “in minutes” analysis of a dataset of 5538 unique viral genomic sequences, for a total of 61. 8 million bp analyzed, with high classification accuracy scores at all levels, from the highest to the lowest taxonomic rank;
- The use of Spearman’s rank correlation analysis to confirm our results and the relatedness of the COVID-19 sequences to the known genera of the family *Coronaviridae* and the known sub-genera of the genus *Betacoronavirus.*

## Materials and methods

The Wuhan seafood market pneumonia virus (COVID-19 virus) isolate Wuhan-Hu-1 complete reference genome of 29903 bp was downloaded from the NCBI database on January 23, 2020. Also, we downloaded all of the available 29 sequences of COVID-19 and the bat *Betacoronavirus RaTG13* from the GISAID platform and two additional sequences (*ba,t-SL-CoVZC45*, and *bat-SL-CoVZXC21*) from the NCBI on January 27, 2019. We downloaded all of the available viral sequences from the Virus-Host DB (14688 sequences available on January 14, 2020). Virus-Host DB covers the sequences from the NCBI RefSeq (release 96, September 9, 2019), and GenBank (release 233.0, August 15, 2019). All sequences shorter than 2000 bp and longer than 50000 bp were ignored to address possible issues arising from sequence length bias.

ML-DSP [51] and MLDSP-GUI [52] were used as the machine learning-based alignment-free methods for complete genome analyses. As MLDSP-GUI is an extension of the ML-DSP methodology, we will refer to the method hereafter as MLDSP-GUI. Each genomic sequence is mapped into its respective genomic signal (a discrete numeric sequence) using a numerical representation. For this study, we use a two-dimensional *k*-mer (oligomers of length *k*) based numerical representation known as Chaos Game Representation (CGR) [53]. The *k*-mer value 7 is used for all the experiments. The value *k* = 7 achieved the highest accuracy scores for the HIV-1 subtype classification [50] and this value could be relevant for other virus related analyses. The magnitude spectra are then calculated by applying Discrete Fourier Transform (DFT) to the genomic signals [51]. A pairwise distance matrix is then computed using the Pearson Correlation Coefficient (PCC) [54] as a distance measure between magnitude spectra. The distance matrix is used to generate the 3D Molecular Distance Maps (MoDMap3D) [55] by applying the classical Multi-Dimensional Scaling (MDS) [56]. MoDMap3D represents an estimation of the relationship among sequences based on the genomic distances between the sequences. The feature vectors are constructed from the columns of the distance matrix and are used as an input to train six supervised-learning based classification models (Linear Discriminant, Linear SVM, Quadratic SVM, Fine KNN, Subspace Discriminant, and Subspace KNN) [51]. A 10-fold cross-validation is used to train, and test the classification models and the average of 10 runs is reported as the classification accuracy. The trained machine learning models are then used to test the COVID-19 sequences. The unweighted pair group method with arithmetic mean (UPGMA) phylogenetic tree is also computed using the pairwise distance matrix.

In this paper, MLDSP-GUI is augmented by a decision tree approach to the supervised machine learning component and a Spearman’s rank correlation coefficient analysis for result validation. The decision tree parallels the taxonomic classification levels, and is necessary so as to minimize the number of calls to the supervised classifier module, as well as to maintain a reasonable number of clusters during each supervised training session. For validation of MLDSP-GUI results using CGR as a numerical representation, we use Spearman’s rank correlation coefficient [57–60], as follows. The frequency of each *k*-mer is calculated in each genome. Due to differences in genome length between species, proportional frequencies are computed by dividing each *k*-mer frequency by the length of the respective sequence. To determine whether there is a correlation between *k*-mer frequencies in COVID-19 and specific taxonomic groups, a Spearman’s rank correlation coefficient test is conducted for *k* = 1 to *k* = 7.

## Results

Table 1 provides the details of three datasets Test-1, Test-2, Test-3a and Test-3b used for analyses with MLDSP-GUI. Each dataset’s composition (clusters with number of sequences), the respective sequence length statistics, and results of MLDSP-GUI after applying 10-fold cross-validation as classification accuracy scores are shown. The classification accuracy scores for all six classification models are shown with their average, see Table 1.

As shown in Table 1, for the first test (Test-1), we organized the dataset of sequences into 12 clusters (11 families, and Riboviria realm). Only the families with at least 100 sequences were considered. The Riboviria cluster contains all families that belong to the realm Riboviria. For the clusters with more than 500 sequences, we selected 500 sequences at random. Our method can handle all of the available 14668 sequences, but using imbalanced clusters, in regard to the number of sequences, can introduce an unwanted bias. After filtering out the sequences, our pre-processed dataset is left with 3273 sequences organized into 12 clusters *(Adenoviridae, Anelloviridae, Caudovirales*, *Geminiviridae*, *Genomoviridae*, *Microviridae*, *Ortervirales*, *Papillomaviridae, Parvoviridae, Polydnaviridae, Polyomaviridae*, and Riboviria). We used MLDSP-GUI with CGR as the numerical representation at *k* = 7. The maximum classification accuracy of 94.9% is obtained using the Quadratic SVM model. The respective MoDMap3D is shown in Figure 1(a). All six classification models trained on 3273 sequences were used to classify (predict the label of) the 29 COVID-19 sequences. All of our machine learning-based models correctly predicted and confirmed the label as Riboviria for all 29 sequences (Table 2).

**Fig 1.**
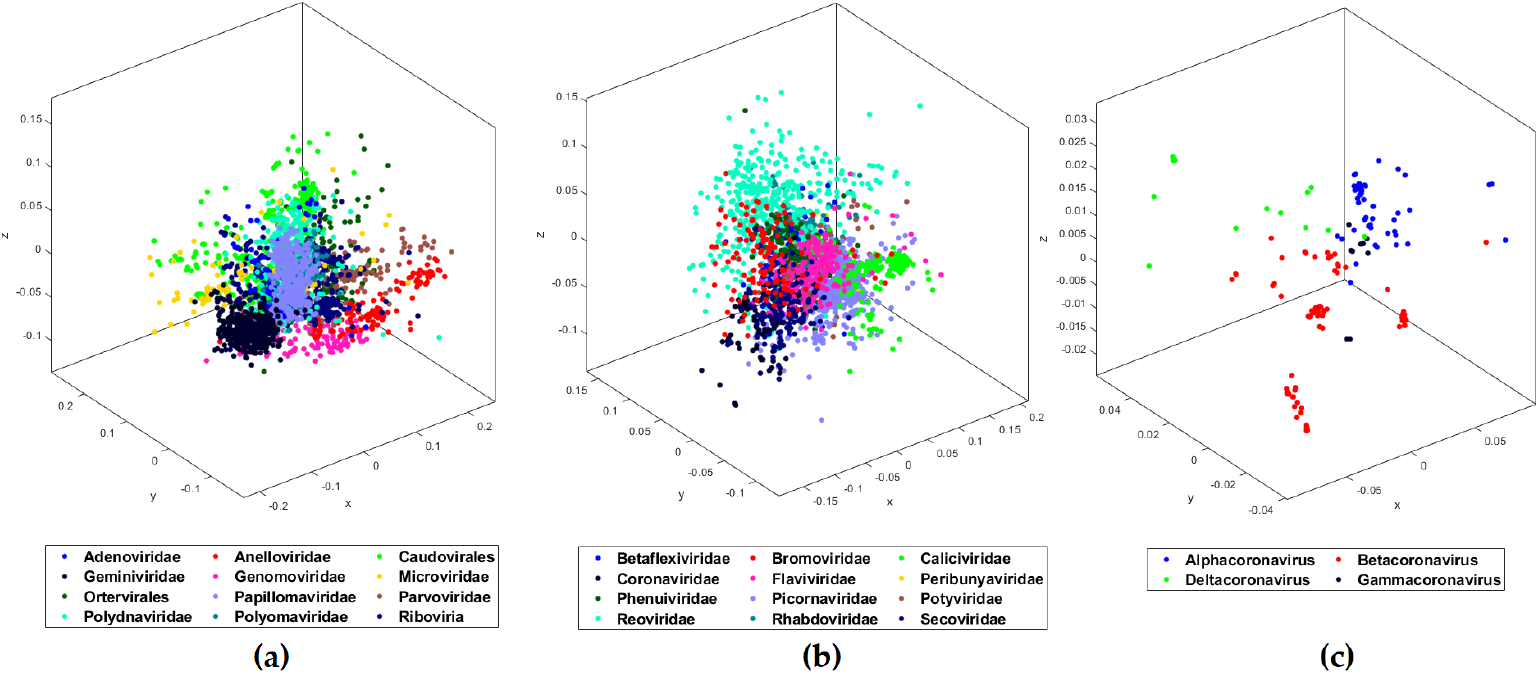
MoDMap3D of (a) 3273 viral sequences from Test-1 representing 11 viral families and realm Riboviria, (b) 2779 viral sequences from Test-2 classifying 12 viral families of realm Riboviria, (c) 208 *Coronaviridae* sequences from Test-3a classified into genera.

Test-1 classified the COVID-19 virus as belonging to the realm Riboviria. The second test (Test-2) is designed to classify COVID-19 among the families of the Riboviria realm. We completed the dataset pre-processing using the same rules as in Test-1 and obtained a dataset of 2779 sequences placed into the 12 families (*Betaflexiviridae, Bromoviridae, Caliciviridae, Coronaviridae, Flaviviridae, Peribunyaviridae, Phenuiviridae, Picornaviridae, Potyviridae, Reoviridae, Rhabdoviridae*, and *Secoviridae*), see Table 1. MLDSP-GUI with CGR at *k* = 7 as the numerical representation was used for the classification of the dataset in Test-2. The maximum classification accuracy of 93.1% is obtained using the Quadratic SVM model. The respective MoDMap3D is shown in Figure 1(b). All six classification models trained on 2779 sequences were used to classify (predict the label of) the 29 COVID-19 sequences. All of our machine learning-based models predicted the label as *Coronaviridae* for all 29 sequences (Table 2) with 100% classification accuracy. Test-2 correctly predicted the family of COVID-19 sequences as *Coronaviridae.* Test-3 performs the genus-level classification.

**Table 1.** Classification accuracy scores of viral sequences at different levels of taxonomy.

| Dataset | Clusters | Number of sequences | Classification model | Classification accuracy (in %) |
| --- | --- | --- | --- | --- |
| Test-1:<br>11 families and Riboviria;<br>3273 sequences;<br>Maximum length: 49973<br>Minimum length: 2002<br>Median length: 7350<br>Mean length: 13173 | *Adenoviridae* | 198 | LinearDiscriminant<br>LinearSVM<br>QuadraticSVM<br>FineKNN<br>SubspaceDiscriminant<br>SubspaceKNN<br>AverageAccuracy | 91.7<br>90.8<br>95<br>93.4<br>87.6<br>93.2<br>92 |
|  | *Anelloviridae* | 126 |  |  |
|  | *Caudovirales* | 500 |  |  |
|  | *Geminiviridae* | 500 |  |  |
|  | *Genomoviridae* | 115 |  |  |
|  | *Microviridae* | 102 |  |  |
|  | *Ortervirales* | 233 |  |  |
|  | *Papillomaviridae* | 369 |  |  |
|  | *Parvoviridae* | 182 |  |  |
|  | *Polydnaviridae* | 304 |  |  |
|  | *Polyomaviridae* | 144 |  |  |
|  | Riboviria | 500 |  |  |
| Test-2:<br>Riboviria families;<br>2779 sequences;<br>Maximum length: 31769<br>Minimum length: 2005<br>Median length: 7488<br>Mean length: 8607 | *Betaflexiviridae* | 121 | LinearDiscriminant<br>LinearSVM<br>QuadraticSVM<br>FineKNN<br>SubspaceDiscriminant<br>SubspaceKNN<br>AverageAccuracy | 91.2<br>89.2<br>93.1<br>90.3<br>89<br>90.4<br>90.5 |
|  | *Bromoviridae* | 122 |  |  |
|  | *Caliciviridae* | 403 |  |  |
|  | *Coronaviridae* | 210 |  |  |
|  | *Flaviviridae* | 222 |  |  |
|  | *Peribunyaviridae* | 166 |  |  |
|  | *Phenuiviridae* | 107 |  |  |
|  | *Picornaviridae* | 437 |  |  |
|  | *Potyviridae* | 196 |  |  |
|  | *Reoviridae* | 470 |  |  |
|  | *Rhabdoviridae* | 192 |  |  |
|  | *Secoviridae* | 133 |  |  |
| Test-3a:<br>*Coronaviridae*;<br>208 sequences;<br>Maximum length: 31769<br>Minimum length: 9580<br>Median length: 29704<br>Mean length: 29256 | *Alphacoronavirus* | 53 | LinearDiscriminant | 98.1 |
|  | *Betacoronavirus* | 126 | LinearSVM | 94.2 |
|  | *Deltacoronavirus* | 20 | QuadraticSVM | 95.2 |
|  | *Gammacoronavirus* | 9 | FineKNN | 95.7 |
|  |  |  | SubspaceDiscriminant | 97.6 |
|  |  |  | SubspaceKNN | 96.2 |
|  |  |  | AverageAccuracy | 96.2 |
| Test-3b:<br>*Coronaviridae*;<br>60 sequences;<br>Maximum length: 31429<br>Minimum length: 25402<br>Median length: 28475<br>Mean length: 28187 | *Alphacoronavirus* | 20 | LinearDiscriminant | 100 |
|  | *Betacoronavirus* | 20 | LinearSVM | 93.3 |
|  | *Deltacoronavirus* | 20 | QuadraticSVM | 93.3 |
|  |  |  | FineKNN | 95 |
|  |  |  | SubspaceDiscriminant | 95 |
|  |  |  | SubspaceKNN | 95 |
|  |  |  | AverageAccuracy | 95.3 |

All classifiers trained on Test-1, Test-2, Test-3a, and Test-3b datasets were used to predict the labels of 29 COVID-19 viral sequences. All classifiers predicted the correct labels for all of the sequences (Riboviria when trained using Test-1, *Coronaviridae* when trained using Test-2, and *Betacoronavirus* when trained using Test-3a and Test-3b).

The third test (Test-3a) is designed to classify the COVID-19 sequences at the genus level. We considered 208 *Coronaviridae* sequences available under four genera *(Alphacoronavirus, Betacoronavirus, Deltacoronavirus, Gammacoronavirus)* (Table 1). MLDSP-GUI with CGR at *k* = 7 as the numerical representation was used for the classification of the dataset in Test-3a. The maximum classification accuracy of 98.1% is obtained using the Linear Discriminant model and the respective MoDMap3D is shown in Figure 1(c). All six classification models trained on 208 sequences were used to classify (predict the label of) the 29 COVID-19 sequences. All of our machine learning-based models predicted the label as *Betacoronavirus* for all 29 sequences (Table 2). To verify that the correct prediction is not an artifact of possible bias because of larger *Betacoronavirus* cluster, we did a secondary Test-3b with cluster size limited to the size of smallest cluster (after removing the *Gammacoronavirus* because it just had 9 sequences). The maximum classification accuracy of 100% is obtained using the Linear Discriminant model for Test-3b. All six classification models trained on 60 sequences were used to classify the 29 COVID-19 sequences. All of our machine learning-based models predicted the label as *Betacoronavirus* for all 29 sequences (Table 2). This secondary test showed that the possible bias is not significant enough to have any impact on the classification performance.

**Table 2.** Predicted taxonomic labels of 29 COVID-19 sequences.

| Training dataset | Testing dataset | Classification models | Prediction accuracy (%) | Predicted label |
| --- | --- | --- | --- | --- |
| Test-1 | 29 COVID-19 Sequences | Linear Discriminant | 100 | Riboviria |
|  |  | Linear SVM | 100 | Riboviria |
|  |  | Quadratic SVM | 100 | Riboviria |
|  |  | Fine KNN | 100 | Riboviria |
|  |  | Subspace Discriminant | 100 | Riboviria |
|  |  | Subspace KNN | 100 | Riboviria |
| Test-2 | 29 COVID-19 Sequences | Linear Discriminant | 100 | *Coronaviridae* |
|  |  | Linear SVM | 100 | *Coronaviridae* |
|  |  | Quadratic SVM | 100 | *Coronaviridae* |
|  |  | Fine KNN | 100 | *Coronaviridae* |
|  |  | Subspace Discriminant | 100 | *Coronaviridae* |
|  |  | Subspace KNN | 100 | *Coronaviridae* |
| Test-3(a\b) | 29 COVID-19 Sequences | Linear Discriminant | 100 | *Betacoronavirus* |
|  |  | Linear SVM | 100 | *Betacoronavirus* |
|  |  | Quadratic SVM | 100 | *Betacoronavirus* |
|  |  | Fine KNN | 100 | *Betacoronavirus* |
|  |  | Subspace Discriminant | 100 | *Betacoronavirus* |
|  |  | Subspace KNN | 100 | *Betacoronavirus* |

Given confirmation that the COVID-19 belongs to the *Betacoronavirus* genus, there now is a question of its origin and relation to the other viruses of the same genus. To examine this question, we preprocessed our dataset from our third test to keep the sub-clusters of the *Betacoronavirus* with at least 10 sequences (Test-4). This gives 124 sequences placed into four clusters (*Embecovirus*, *Merbecovirus*, *Nobecovirus*, *Sarbecovirus)* (Table 3). The maximum classification accuracy of 98.4% with CGR at *k* = 7 as the numerical representation is obtained using the Quadratic SVM model. The respective MoDMap3D is shown in Figure 2(a). All six classifiers trained on 124 sequences predicted the label as *Sarbecovirus*, when used to predict the labels of 29 COVID-19 sequences. For Test-5, we added COVID-19 with 29 sequences as the fifth cluster, see Table 3. The maximum classification accuracy of 98.7% with CGR at *k* = 7 as the numerical representation is obtained using the Subspace Discriminant model. The respective MoDMap3D is shown in Figure 2(b). In the MoDMap3D plot from Test-5, COVID-19 sequences are placed in a single distinct cluster, see Figure 2(b). As visually suggested by the MoDMap3D (Figure 2(b)), the average inter-cluster distances confirm that the COVID-19 sequences are closest to the *Sarbecovirus* (average distance 0.0556), followed by *Merbecovirus* (0.0746), *Embecovirus* (0.0914), and *Nobecovirus* (0.0916). The three closest sequences based on the average distances from all COVID-19 sequences are *RaTG13* (0.0203), *bat-SL-CoVZCĄ5* (0.0418), and *bat-SL-CoVZXC21* (0.0428).

**Fig 2.**
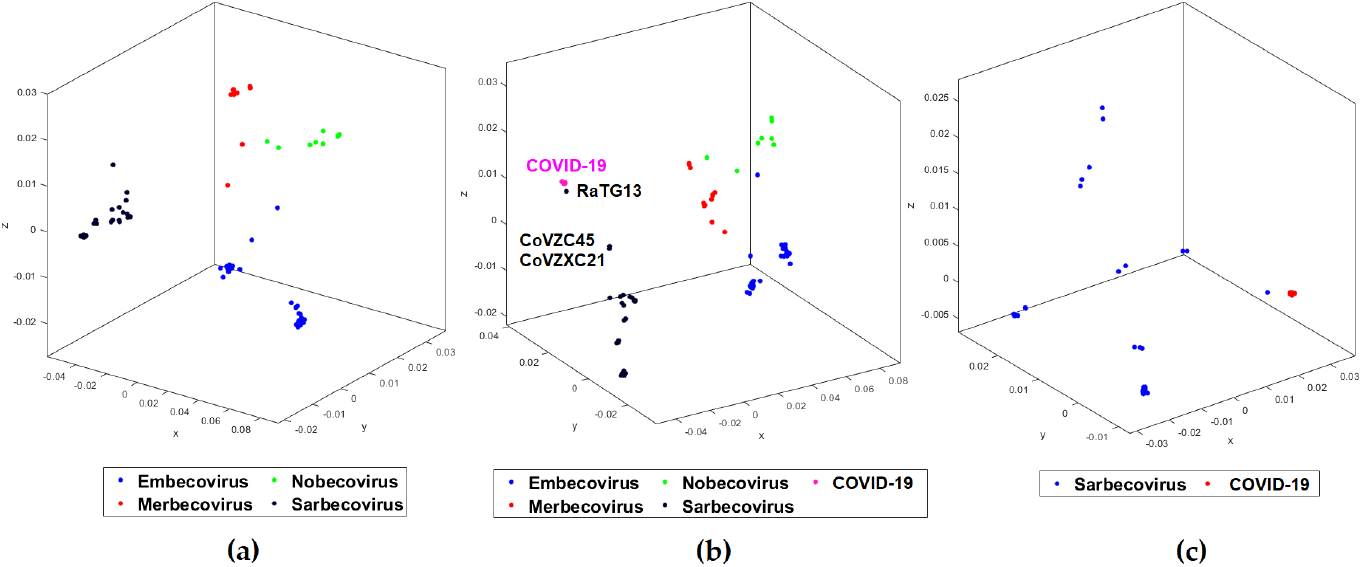
MoDMap3D of (a) 124 *Betacoronavirus* sequences from Test-4 classified into sub-genera, (b) 153 viral sequences from Test-5 classified into 4 sub-genera and COVID-19, (c) 76 viral sequences from Test 6 classified into *Sarbecovirus* and COVID-19.

**Table 3.** Genus to sub-genus classification accuracy scores of *Betacoronavirus*.

| Dataset | Clusters | Number of sequences | Classification model | Classification accuracy (in %) |
| --- | --- | --- | --- | --- |
| Test-4:<br>*Betacoronavirus*;<br>124 sequences;<br>Maximum length: 31526<br>Minimum length: 29107<br>Median length: 30155<br>Mean length: 30300 | *Embecovirus*<br>*Merbecovirus*<br>*Nobecovirus*<br>*Sarbecovirus* | 49<br>18<br>10<br>47 | LinearDiscriminant | 97.6 |
|  |  |  | LinearSVM | 98.4 |
|  |  |  | QuadraticSVM | 98.4 |
|  |  |  | FineKNN | 97.6 |
|  |  |  | SubspaceDiscriminant | 98.4 |
|  |  |  | SubspaceKNN | 97.2 |
|  |  |  | AverageAccuracy | 97.6 |
| Test-5:<br>*Betacoronavirus* and<br>COVID-19;<br>153 sequences;<br>Maximum length: 31526<br>Minimum length: 29107<br>Median length: 29891<br>Mean length: 30217 | *Embecovirus*<br>*Merbecovirus*<br>*Nobecovirus*<br>*Sarbecovirus*<br>COVID-19 | 49<br>18<br>10<br>47<br>29 | LinearDiscriminant | 98.6 |
|  |  |  | LinearSVM | 97.4 |
|  |  |  | QuadraticSVM | 97.4 |
|  |  |  | FineKNN | 97.4 |
|  |  |  | SubspaceDiscriminant | 98.7 |
|  |  |  | SubspaceKNN | 96.1 |
|  |  |  | AverageAccuracy | 97.5 |
| Test-6:<br>*Sarbecovirus* and<br>COVID-19;<br>76 sequences;<br>Maximum length: 30309<br>Minimum length: 29452<br>Median length: 29748<br>Mean length: 29772 | *Sarbecovirus*<br>COVID-19 | 47<br>29 | LinearDiscriminant | 100 |
|  |  |  | LinearSVM | 100 |
|  |  |  | QuadraticSVM | 100 |
|  |  |  | FineKNN | 100 |
|  |  |  | SubspaceDiscriminant | 100 |
|  |  |  | SubspaceKNN | 100 |
|  |  |  | AverageAccuracy | 100 |

For Test-6, we classified *Sarbecovirus* (47 sequences) and COVID-19 (29 sequences) clusters and achieved separation of the two clusters visually apparent in the MoDMap3D, see Figure 2(c). Quantitatively, using 10-fold cross-validation, all six of our classifiers report 100% classification accuracy. We generated a phylogenetic tree based on all pairwise distances for the dataset in Test-6 that shows the separation of the two clusters and relationships within the clusters (Figure 3). As observed in Test-5, the phylogenetic tree shows that the COVID-19 sequences are closer to the bat *Betacoronavirus RaTG13* sequence collected from a bat host.

**Fig 3.**
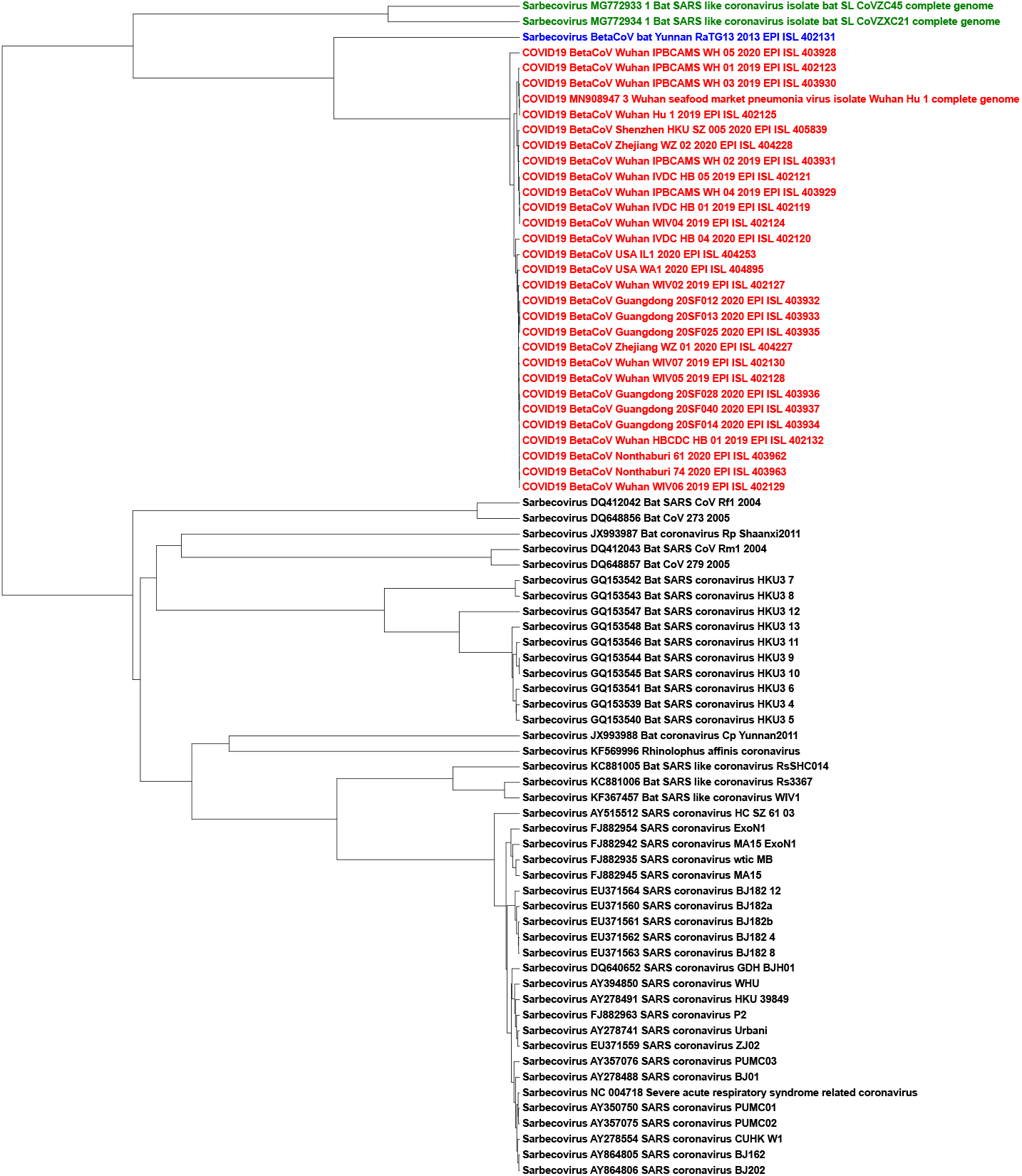
The UPGMA phylogenetic tree using the Pearson Correlation Coefficient generated pairwise distance matrix shows COVID-19 (Red) sequences proximal to the bat *Betacoronavirus* RaTG13 (Blue) and bat SARS-like coronaviruses ZC45/ZXC21 (Green) in a distinct lineage from the rest of *Sarbecovirus* sequences

Figure 4 shows the Chaos Game Representation (CGR) plots of different sequences from the four different genera (*Alphacoronavirus*, *Betacoronavirus*, *Deltacoronavirus*, *Gammacoronavirus)* of the family *Coronaviridae.* The CGR plots visually suggests and the pairwise distances confirm that the genomic signature of the COVID-19 Wuhan-Hu-1 (Figure 4(a)) is closer to the genomic signature of the *BetaCov-RaTG13* (Figure 4(b); distance: 0.0204), followed by the genomic signatures of *bat-SL-CoVZCĄδ* (Figure 4(c); distance: 0.0417), *bat-SL-CoVZXC21(Figure 4(d)*; distance: 0.0428), *Alphacoronavirus /DQ811787 PRCV ISU*-1 (Figure 4(e); distance: 0.0672), *Gammacoronavirus /* Infectious bronchitis virus NGA /A116E*7/2006/FN*430415 (Figure 4(f); distance: 0.0791), and *Deltacoronavirus /* PDCoV / USA / Illinois121 /2014/KJ481931 (Figure 4(g); distance: 0.0851).

**Fig 4.**
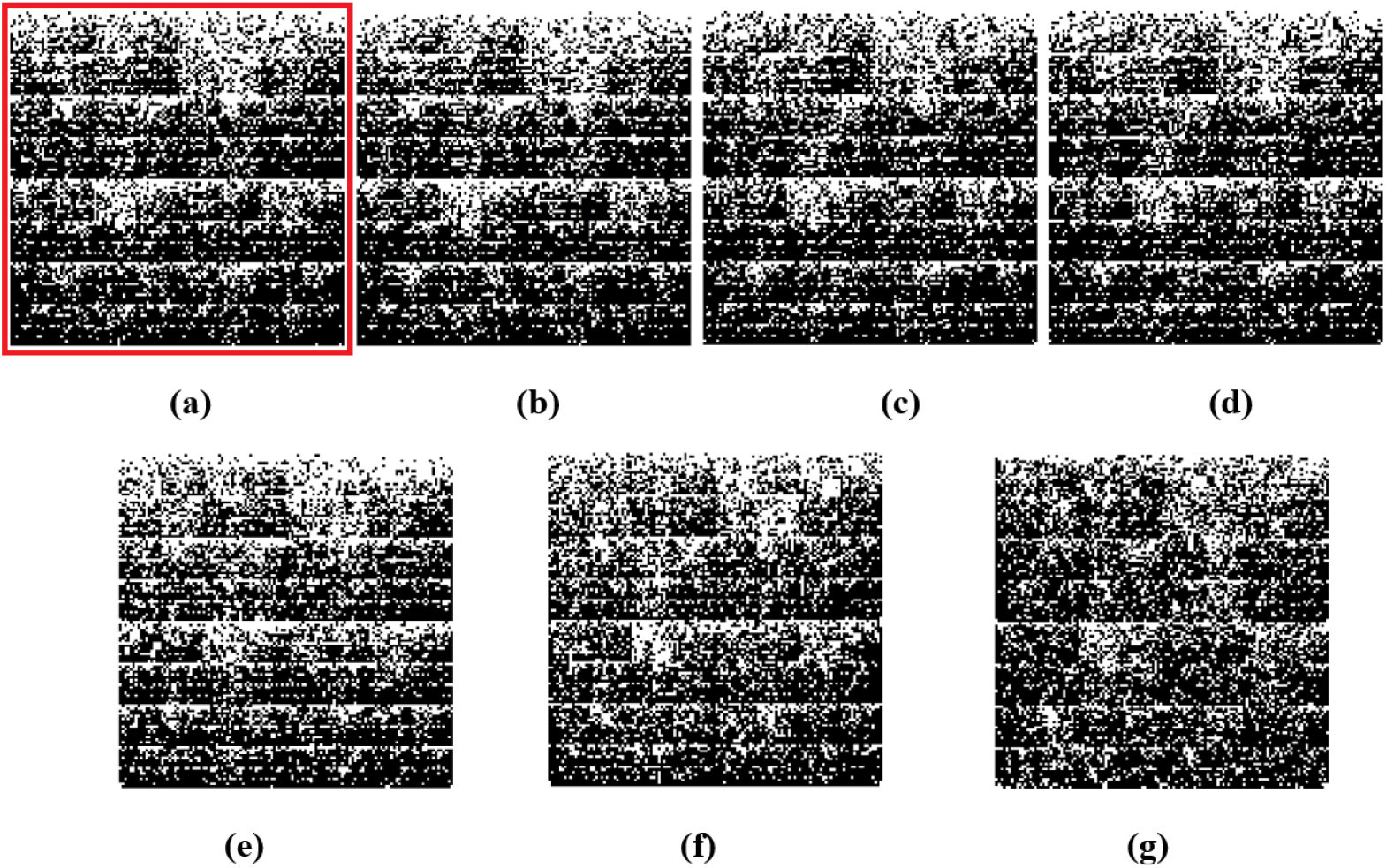
Chaos Game Representation (CGR) plots at *k* = 7 of (a) COVID-19 / Wuhan seafood market pneumonia virus isolate Wuhan-Hu-1/MN908947.3, (b) *Betacoronavirus /* CoV / Bat / Yunnan / RaTG13 /EPI_ISL_402131, (c) *Betacoronavirus /* Bat SARS-like coronavirus isolate bat-SL-CoVZC45 /MG772933.1, (d) *Betacoronavirus /* Bat SARS-like coronavirus isolate bat-SL-CoVZXC21 /MG772934.1, (e) *Alphacoronavirus /DQ811787 PRCV ISU*-1, (f) *Gammacoronavirus /* Infectious bronchitis virus NGA /A116E7/2006/FN430415, and (g) *Deltacoronavirus /* PDCoV / USA / Illinois121 /2014/KJ481931. Chaos plot vertices are assigned top left Cytosine, top right Guanine, bottom left Adenine and bottom right Thymine.

The Spearman’s rank correlation coefficient tests were used to further confirm the ML-DSP findings. The first test in Figure 5 shows the COVID-19 being compared to the four genera; *Alphacoronavirus*, *Betacoronavirus*, *Gammacoronavirus* and *Deltacoronavirus.* The COVID-19 showed the highest *k*-mer frequency correlation to *Betacoronavirus* at *k* = 7 (Table 4), which is consistent with the ML-DSP results in Test-3 (Table 2). The COVID-19 was then compared to all sub-genera within the *Betacoronavirus* genus: *Embecovirus, Merbecovirus, Nobecovirs* and *Sarbecovirus* seen in Figure 6. The Spearman’s rank test was again consistent with the ML-DSP results seen in Table 3, as the *k*-mer frequencies at *k* = 7 showed the highest correlation to the sub-genus *Sarbecovirus* (Table 4). These tests confirm the findings in ML-DSP and are consistent with the COVID-19 virus as part of the sub-genus *Sarbecovirus.*

**Fig 5.**
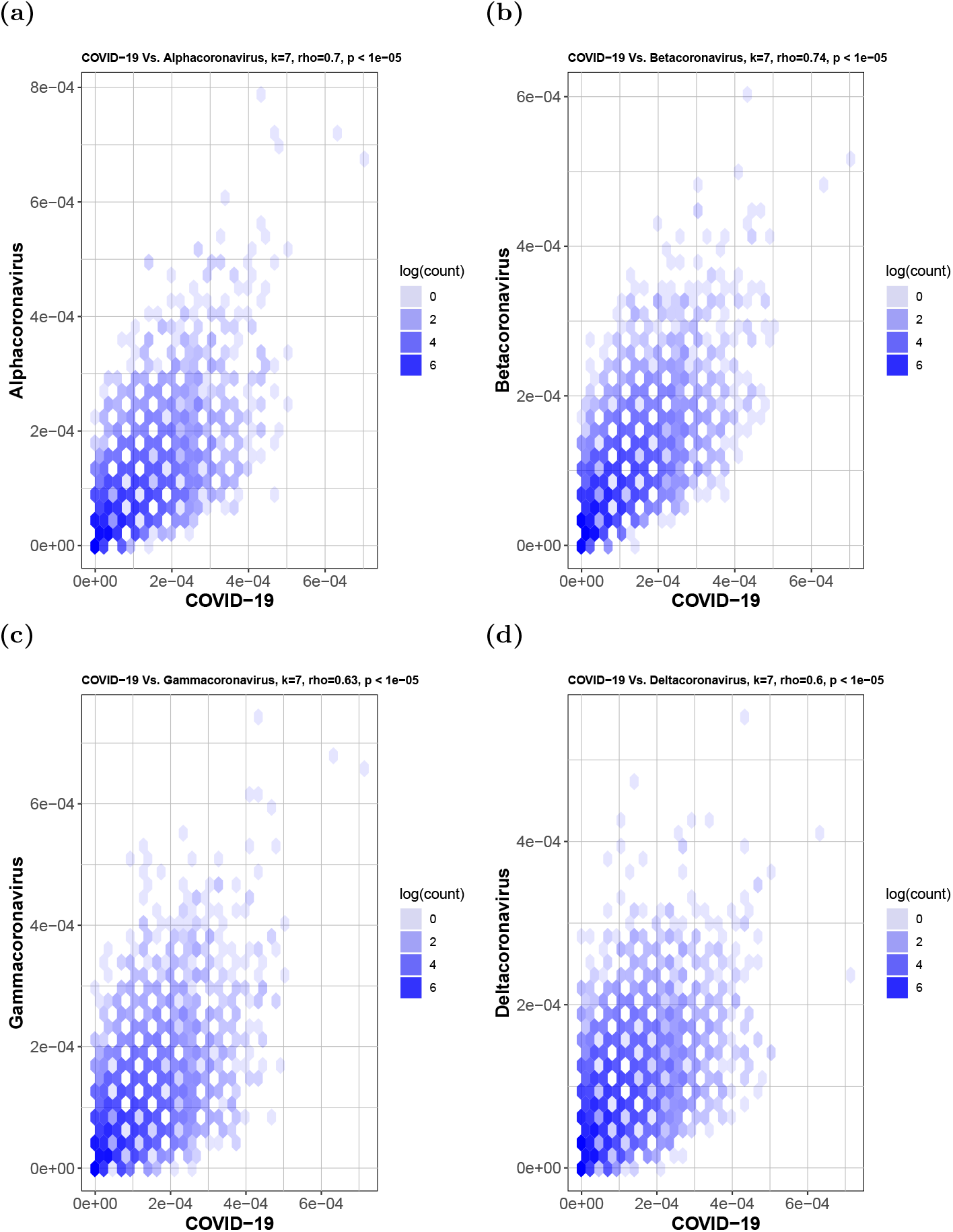
Hexbin scatterplots of the proportional *k*-mer (*k* = 7) frequencies of the COVID-19 sequences versus the four genera: (a) *Alphacoronavirus, ρ* = 0.7; (b) *Betacoronavirus, ρ* = 0.74; (c) *Gammacoronavirus, ρ* = 0.63 and (d) *Deltacoronavirus, ρ* = 0.6. The color of each hexagonal bin in the plot represents the number of points (in natural logarithm scale) overlapping at that position. All *ρ* values resulted in *p*-values < 10^-5^ for the correlation test. By visually inspecting each hexbin scatterplot, the degree of correlation is displayed by the variation in spread between the points. Hexagonal points that are closer together and less dispersed as seen in (b) are more strongly correlated and have less deviation.

**Fig 6.**
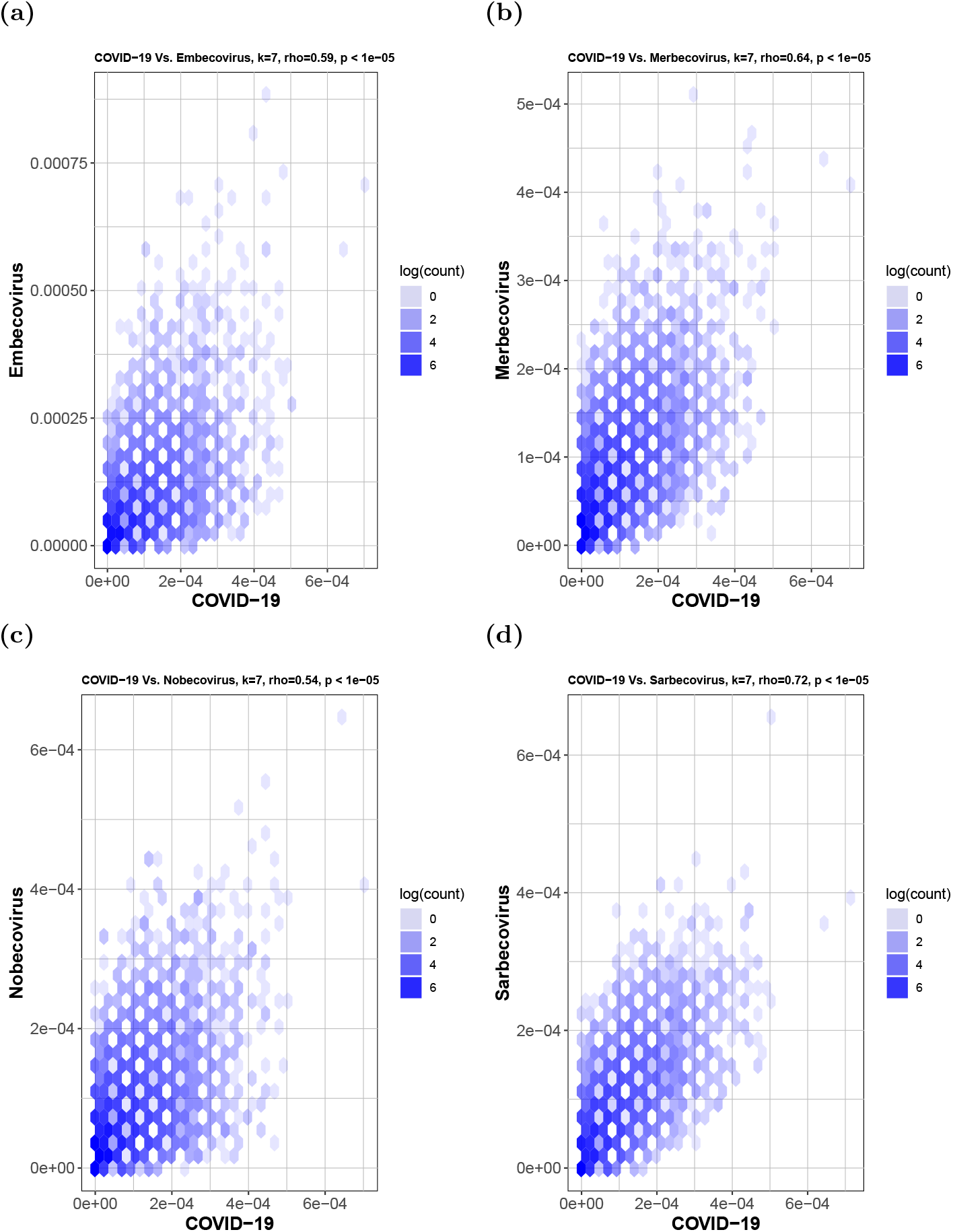
Hexbin scatterplots of the proportional *k*-mer (*k* = 7) frequencies of the COVID-19 sequences versus the four sub-genera: (a) *Embecovirus, ρ* = 0.59; (b) *Merbecovirus, ρ* = 0.64; (c) *Nobecovirus, ρ* = 0.54 and (d) *Sarbecovirus, ρ* = 0.72. The color of each hexagonal bin in the plot represents the number of points (in natural logarithm scale) overlapping at that position. All *ρ* values resulted in *p*-values < 10^-5^ for the correlation test. By visually inspecting each hexbin scatterplot, the degree of correlation is displayed by the variation in spread between the points. Hexagonal points that are closer together and less dispersed as seen in (d) are more strongly correlated and have less deviation.

**Table 4.**
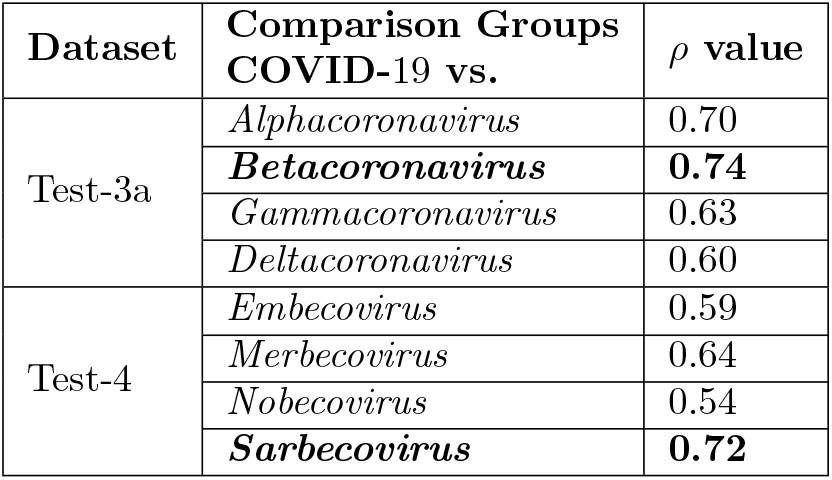
Spearman’s rank correlation coefficient (*ρ*) values from Figure 5 and 6, for which all *p*-values < 10^-5^. The strongest correlation value was found between *Betacoronavirus* and *Sarbecovirus* when using the data sets from Test 3a from Table 2 and Test 4 from Table 3, respectively.

## Discussion

Prior work elucidating the evolutionary history of the Wuhan COVID-19 virus had suggested an origin from bats prior to zoonotic transmission [12,34,36,39,42,61]. Most early cases of individuals infected with COVID-19 had contact with the Huanan South China Seafood Market [26–31]. Human-to-human transmission is confirmed, further highlighting the need for continued intervention [34,61–63]. Still, the early COVID-19 genomes that have been sequenced and uploaded are over 99% similar, suggesting these infections result from a recent cross-species event [12,31,41].

These prior analyses relied upon alignment-based methods to identify relationships between COVID-19 and other coronaviruses with nucleotide and amino acid sequence similarities. When analyzing the conserved replicase domains of ORF1ab for coronavirus species classification, nearly 94% of amino acid residues were identical to SARS-CoV, yet overall genome similarity was only around 70%, confirming that COVID-19 was genetically different [63]. Within the RdRp region, it was found that another bat coronavirus, *RaTG13*, was the closest relative to COVID-19 and formed a distinct lineage from other bat SARS-like coronaviruses [39,41]. Other groups found that two bat SARS-like coronaviruses, *bat-SL-CoVZCĄδ* and *bat-SL-CoVZXC21*, were also closely related to COVID-19 [12,34–38]. There is a consensus that these three bat viruses are most similar to COVID-19, however, whether or not COVID-19 arose from a recombination event is still unknown [39–41].

Regardless of the stance on recombination, current consensus holds that the hypothesis of COVID-19 originating from bats is highly likely. Bats have been identified as a reservoir of mammalian viruses and cross-species transmission to other mammals, including humans [4, 7, 8, 10, 13, 64–66]. Prior to intermediary cross-species infection, the coronaviruses SARS-CoV and MERS-CoV were also thought to have originated in bats [24,25,35,68,69]. Many novel SARS-like coronaviruses have been discovered in bats across China, and even in European, African and other Asian countries [35, 70–76]. With widespread geographic coverage, SARS-like coronaviruses have likely been present in bats for a long period of time and novel strains of these coronaviruses can arise through recombination [4]. Whether or not COVID-19 was transmitted directly from bats, or from intermediary hosts, is still unknown, and will require identification of COVID-19 in species other than humans, notably from the wet market and surrounding area it is thought to have originated from [30]. While bats have been reported to have been sold at the Huanan market, at this time, it is still unknown if there were intermediary hosts involved prior to transmission to humans [27,31,34,40,77]. Snakes had been proposed as an intermediary host for COVID-19 based on relative synonymous codon usage bias studies between viruses and their hosts [40], however, this claim has been disputed [78]. China CDC released information about environmental sampling in the market and indicated that 33 of 585 samples had evidence of COVID-19, with 31 of these positive samples taken from the location where wildlife booths were concentrated, suggesting possible wildlife origin [79,80]. Detection of SARS-CoV in Himalyan palm civets and horseshoe bats identified 29 nucleotide sequences that helped trace the origins of SARS-CoV isolates in humans to these intermediary species [13, 24, 39, 76]. Sampling additional animals at the market and wildlife in the surrounding area may help elucidate whether intermediary species were involved or not, as was possible with the SARS-CoV.

Viral outbreaks like COVID-19 demand timely analysis of genomic sequences to guide the research in the right direction. This problem being time-sensitive requires quick sequence similarity comparison against thousands of known sequences to narrow down the candidates of possible origin. Alignment-based methods are known to be time-consuming and can be challenging in cases where homologous sequence continuity cannot be ensured. It is challenging (and sometimes impossible) for alignment-based methods to compare a large number of sequences that are too different in their composition. Alignment-free methods have been used successfully in the past to address the limitations of the alignment-based methods [49–52]. The alignment-free approach is quick and can handle a large number of sequences. Moreover, even the sequences coming from different regions with different compositions can be easily compared quantitatively, with equally meaningful results as when comparing homologous/similar sequences. We use MLDSP-GUI (a variant of MLDSP with additional features), a machine learning-based alignment-free method successfully used in the past for sequence comparisons and analyses [51]. The main advantage alignment-free methodology offers is the ability to analyze large datasets rapidly. In this study we confirm the taxonomy of COVID-19 and, more generally, propose a method to efficiently analyze and classify a novel unclassified DNA sequence against the background of a large dataset. We namely use a “decision tree” approach (paralleling taxonomic ranks), and start with the highest taxonomic level, train the classification models on the available complete genomes, test the novel unknown sequences to predict the label among the labels of the training dataset, move to the next taxonomic level, and repeat the whole process down to the lowest taxonomic label.

Test-1 starts at the highest available level and classifies the viral sequences to the 11 families and Riboviria realm (Table 1). There is only one realm available in the viral taxonomy, so all of the families that belong to the realm Riboviria are placed into a single cluster and a random collection of 500 sequences are selected. No realm is defined for the remaining 11 families. The objective is to train the classification models with the known viral genomes and then predict the labels of the COVID-19 virus sequences. The maximum classification accuracy score of 95% was obtained using the Quadratic SVM model. This test demonstrates that MLDSP-GUI can distinguish between different viral families. The trained models are then used to predict the labels of 29 COVID-19 sequences. As expected, all classification models correctly predict that the COVID-19 sequences belong to the Riboviria realm, see Table 2. Test-2 is composed of 12 families from the Riboviria, see Table 1, and the goal is to test if MLDSP-GUI is sensitive enough to classify the sequences at the next lower taxonomic level. It should be noted that as we move down the taxonomic levels, sequences become much more similar to one another and the classification problem becomes challenging. MLDSP-GUI is still able to distinguish between the sequences within the Riboviria realm with a maximum classification accuracy of 91.1% obtained using the Linear Discriminant classification model. When COVID-19 sequences are tested using the models trained on Test-2, all of the models correctly predict the COVID-19 sequences as *Coronaviridae* (Table 2). Test-3a moves down another taxonomic level and classifies the *Coronaviridae* family to four genera *(Alphacoronavirus, Betacoronavirus, Deltacoronavirus, Gammacoronavirus)*, see Table 1. MLDSP-GUI distinguishes sequences at the genus level with a maximum classification accuracy score of 98%, obtained using the Linear Discriminant model. This is a very high accuracy rate considering that no alignment is involved and the sequences are very similar. All trained classification models correctly predict the COVID-19 as *Betacoronavirus*, see Table 2. Test-3a has *Betacoronavirus* as the largest cluster and it can be argued that the higher accuracy could be a result of this bias. To avoid bias, we did an additional test removing the smallest cluster *Gammacoronavirus* and limiting the size of remaining three clusters to the size of the cluster with the minimum number of sequences i.e. 20 with Test-3b. MLDSP-GUI obtains 100% classification accuracy for this additional test and still predicts all of the COVID-19 sequences as *Betacoronavirus.* These tests confirm that the COVID-19 are from the genus *Betacoronavirus.*

Sequences become very similar at lower taxonomic levels (sub-genera and species). Test-4, Test-5, and Test-6 investigate within the genus *Betacoronavirus* for sub-genus classification. Test-4 is designed to classify *Betacoronavirus* into the four sub-genera *(Embecovirus, Merbecovirus, Nobecovirus, Sarbecovirus)*, see Table 3. MLDSP-GUI distinguishes sequences at the sub-genus level with a maximum classification accuracy score of 98.4%, obtained using the Quadratic SVM model. All of the classification models trained on the dataset in Test-4 predicted the label of all 29 COVID-19 sequences as *Sarbecovirus*. This suggests substantial similarity between COVID-19 and the *Sarbecovirus* sequences. Test-5 and Test-6 (see Table 3) are designed to verify that COVID-19 sequences can be differentiated from the known species in the *Betacoronavirus* genus. MLDSP-GUI achieved a maximum classification score of 98.7% for Test-5 and 100% for Test-6 using Subspace Discriminant classification model. This shows that although COVID-19 and *Sarbecovirus* are closer on the basis of genomic similarity (Test-4), they are still distinguishable from known species. Therefore, these results suggest that COVID-19 may represent a genetically distinct species of *Sarbecovirus.* All COVID-19 virues are visually seen in MoDMap3D generated from Test-5 (see Figure 2(b)) as a closely packed cluster and it supports a fact that there is 99% similarity among these sequences [12,31]. The MoDMap3D generated from the Test-5 (Figure 2(b)) visually suggests and the average distances from COVID-19 sequences to all other sequences confirm that the COVID-19 sequences are most proximal to the *RaTG13* (distance: 0.0203), followed by the *bat-SL-CoVZCĄ5* (0.0418), and *bat-SL-CoVZX21* (0.0428). To confirm this proximity, a UPGMA phylogenetic tree is computed from the PCC-based pairwise distance matrix of sequences in Test-6, see Figure 3. The phylogenetic tree placed the *RaTG13* sequence closest to the COVID-19 sequences, followed by the *bat-SL-CoVZCĄ.5* and *bat-SL-CoVZX21* sequences. This closer proximity represents the smaller genetic distances between these sequences and aligns with the visual sequence relationships shown in the MoDMap3D of Figure 2(b).

We further confirm our results regarding the closeness of COVID-19 with the sequences from the *Betacoronavirus* genus (especially sub-genus *Sarbecovirus)* by a quantitative analysis based on the Spearman’s rank correlation coefficient tests. Spearman’s rank correlation coefficient [57–60] tests were applied to the frequencies of oligonucleotide segments, adjusting for the total number of segments, to measure the degree and statistical significance of correlation between two sets of genomic sequences. Spearman’s ρ value provides the degree of correlation between the two groups and their *k*-mer frequencies. The COVID-19 virus was compared to all genera under the *Coronaviridae* family and the *k*-mer frequencies showed the strongest correlation to the genus *Betacoronavirus*, and more specifically *Sarbecovirus.* The Spearman’s rank tests corroborate that the COVID-19 virus is part of the *Sarbecovirus* sub-genus, as shown by CGR and ML-DSP. When analyzing sub-genera, it could be hard to classify at lower *k* values due to the short oligonucleotide frequencies not capturing enough information to highlight the distinctions. Therefore despite the Spearman’s rank correlation coefficient providing results for *k* = 1 to *k* = 7, the higher *k*-mer lengths provided more accurate results, and *k* = 7 was used.

Attributes of the COVID-19 genomic signature are consistent with previously reported mechanisms of innate immunity operating in bats as a host reservoir for coronaviruses. Vertebrate genomes are known to have an under-representation of CG dinucleotides in their genomes, otherwise known as CG suppression [81,82]. This feature is thought to have been due to the accumulation of spontaneous deamination mutations of methyl-cytosines over time [81]. As viruses are obligate parasites, evolution of viral genomes is intimately tied to the biology of their hosts [83]. As host cells develop strategies such as RNA interference and restriction-modification systems to prevent and limit viral infections, viruses will continue to counteract these strategies [82–84]. Dinucleotide composition and biases are pervasive across the genome and make up a part of the organism’s genomic signature [83]. These host genomes have evolutionary pressures that shape the host genomic signature, such as the pressure to eliminate CG dinucleotides within protein coding genes in humans [82]. Viral genomes have been shown to mimic the same patterns of the hosts, including single-stranded positive-sense RNA viruses, which suggests that many RNA viruses can evolve to mimic the same features of their host’s genes and genomic signature [81–85]. As genomic composition, specifically in mRNA, can be used as a way of discriminating self vs non-self RNA, the viral genomes are likely shaped by the same pressures that influence the host genome [82]. One such pressure on DNA and RNA is the APOBEC family of enzymes, members of which are known to cause G to A mutations [85–87]. While these enzymes primarily work on DNA, it has been demonstrated that these enzymes can also target RNA viral genomes [86]. The APOBEC enzymes therefore have RNA editing capability and may help contribute to the innate defence system against various RNA viruses [85]. This could therefore have a direct impact on the genomic signature of RNA viruses. Additional mammalian mechanisms for inhibiting viral RNA have been highlighted for retroviruses with the actions of zinc-finger antiviral protein (ZAP) [81]. ZAP targets CG dinucleotide sequences, and in vertebrate host cells with the CG suppression in host genomes, this can serve as a mechanism for the distinction of self vs non-self RNA and inhibitory consequences [81]. Coronaviruses have A/U rich and C/G poor genomes, which over time may have been, in part, a product of cytidine deamination and selection against CG dinucleotides [88–90]. This is consistent with the fact that bats serve as a reservoir for many coronaviruses and that bats have been observed to have some of the largest and most diverse arrays of APOBEC genes in mammals [66, 67]. The Spearman’s rank correlation data and the patterns observed in the CGR images from Figure 4, of the coronavirus genomes, including COVID-19 identify patterns such as CG underepresentation, also present in vertebrate and, importantly, bat host genomes.

With human-to-human transmission confirmed and concerns for possible asymptomatic transmission, there is a strong need for continued intervention to prevent the spread of the virus [33,34,61–63]. Due to the high amino acid similarities between COVID-19 and SARS-CoV main protease essential for viral replication and processing, anticoronaviral drugs targeting this protein and other potential drugs have been identified using virtual docking to the protease for treatment of COVID-19 [29,44,45,91–94]. The human ACE2 receptor has also been identified as the potential receptor for COVID-19 and represents a potential target for treatment [42,43].

MLDSP-GUI is an ultra-fast, alignment-free method as is evidenced by the time-performance of MLDSP-GUI for Test-1 to Test-6 given in Figure 7. MLDSP-GUI took just 10.55 seconds to compute a pairwise distance matrix (including reading sequences, computing magnitude spectra using DFT, and calculating the distance matrix using PCC combined) for the Test-1 (largest dataset used in this study with 3273 complete genomes). All of the tests combined (Test-1 to Test-6) are doable in under 10 minutes including the computationally heavy 10-fold cross-validation, and testing of 29 COVID-19 sequences.

**Fig 7.**
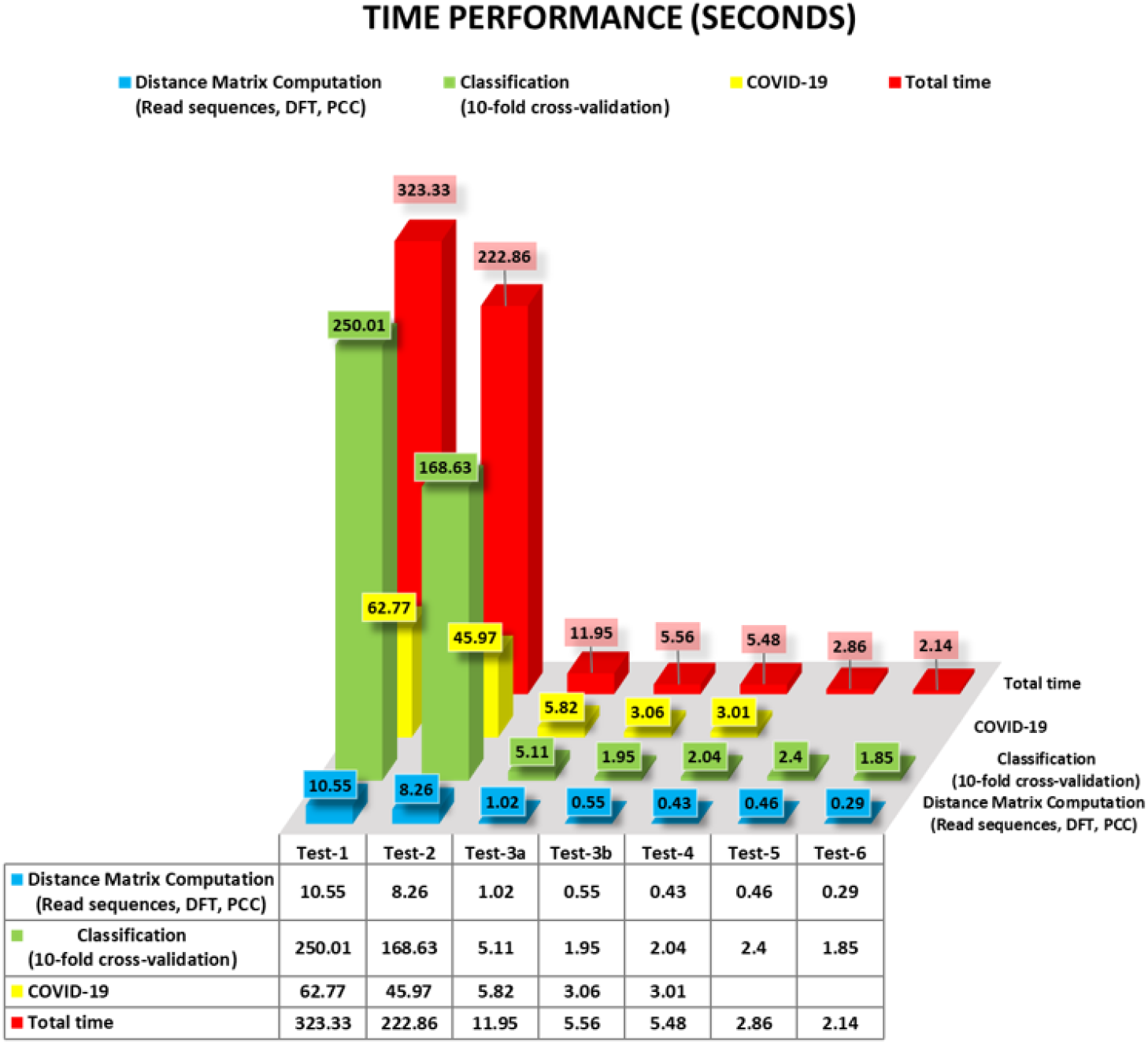
Time performance of MLDSP-GUI for Test1 to Test-6 (in seconds).

The results of our machine learning-based alignment-free analyses using MLDSP-GUI support the hypothesis of a bat origin for COVID-19 and classify COVID-19 as sub-genus *Sarbecovirus*, within *Betacoronavirus*.

## Conclusion

This study provides an alignment-free method based on intrinsic genomic signatures that can deliver highly-accurate real-time taxonomic predictions of yet unclassified new sequences, *ab initio*, using raw DNA sequence data alone and without the need for gene or genome annotation. We use this method to provide evidence for the taxonomic classification of COVID-19 as *Sarbecovirus*, within *Betacoronavirus*, as well as quantitative evidence supporting a bat origin hypothesis. Our results are obtained through a comprehensive analysis of over 5000 unique viral sequences, through an alignment-free analysis of their two-dimensional genomic signatures, combined with a “decision tree” use of supervised machine learning and confirmed by Spearman’s rank correlation coefficient analyses. This study suggests that such alignment-free approaches to comparative genomics can be used to complement alignment-based approaches when timely taxonomic classification is of the essence, such as at critical periods during novel viral outbreaks.

## Supporting information

S1 Table. Spearman’s rank correlation coefficient (*ρ*) value for *k* =1 to *k* = 7.

S2 Table. Accession IDs of the Sequences downloaded from the GISAID.

S3 Table. Accession IDs of sequences used in Test-1 to Test-6.

## Acknowledgments

The authors are appreciative of the review of a manuscript draft by Hailie Pavanel.

## References

1. Enjuanes L, Brian D, Cavanagh D, Holmes K, Lai MMC, Laude H, et al. Coronaviridae. In: Regenmortel MV, Fauquet CM, Bishop DHL, Carstens EB, Estes MK, Lemon SM, et al., editors. Virus Taxonomy. Seventh Report of the International Committee on Taxonomy of Viruses, Academic Press; 2000. pp. 835–849.

2. Weiss SR, Navas-Martin S. Coronavirus Pathogenesis and the Emerging Pathogen Severe Acute Respiratory Syndrome Coronavirus. Microbiol. Mol. Biol. 2005; Rev. 69: 635–664.

3. Su S, Wong G, Shi W, Liu J, Lai ACK, Zhou J, et al. Epidemiology, Genetic Recombination, and Pathogenesis of Coronaviruses. Trends in Microbiology. 2016; 24: 490–502.

4. Cui J, Li F, Shi ZL. Origin and evolution of pathogenic coronaviruses. Nature Reviews Microbiology. 2019; 17: 181–5192.

5. Schoeman D, Fielding BC. Coronavirus envelope protein: Current knowledge. Virology Journal. 2019; 16.

6. de Groot RJ, Baker SC, Baric R, Enjuanes L, Gorbalenya AE, Holmes KV, et al. Family Coronaviridae. In: King AMQ, Adams MJ, Carstens EB, Lefkowitz EJ, editors. Virus taxonomy. Ninth report of the international committee on taxonomy of viruses, Elsevier Academic Press; 2012. pp. 806–828.

7. Woo PCY, Lau SKP, Huang Y, Yuen KY. Coronavirus diversity, phylogeny and interspecies jumping. Experimental Biology and Medicine. 2009; 234: 1117–1127.

8. Wertheim JO, Chu DKW, Peiris JSM, Kosakovsky Pond SL, Poon LLM. A Case for the Ancient Origin of Coronaviruses. J. Virol. 2013; 87: 7039–7045.

9. Luk HKH, Li X, Fung J, Lau SKP, Woo PCY. Molecular epidemiology, evolution and phylogeny of SARS coronavirus. Infection, Genetics and Evolution. 2019; 71: 21–30.

10. Vijaykrishna D, Smith GJD, Zhang JX, Peiris JSM, Chen H, Guan Y. Evolutionary Insights into the Ecology of Coronaviruses. J. Virol. 2007; 81: 4012–4020.

11. Lau SK, Li KS, Tsang AK, Shek CT, Wang M, Choi GK, et al. Recent Transmission of a Novel Alphacoronavirus, Bat Coronavirus HKU10, from Leschenault’s Rousettes to Pomona Leaf-Nosed Bats: First Evidence of Interspecies Transmission of Coronavirus between Bats of Different Suborders. J. Virol. 2012; 86: 11906–11918.

12. Lu R, Zhao X, Li J, Niu P, Yang B, Wu H, et al. Genomic characterisation and epidemiology of 2019 novel coronavirus: implications for virus origins and receptor binding. Lancet. 2020; doi:10.1016/S0140-6736(20)30251-8.

13. Li W, Shi Z, Yu M, Ren W, Smith C, Epstein JH, et al. Bats are natural reservoirs of SARS-like coronaviruses. Science. 2005; 310: 676–679.

14. Duffy S, Shackelton LA, Holmes EC. Rates of evolutionary change in viruses: Patterns and determinants. Nature Reviews Genetics. 2008; 9: 267–276.

15. Jenkins GM, Rambaut A, Pybus OG, Holmes EC. Rates of molecular evolution in RNA viruses: A quantitative phylogenetic analysis. J. Mol. Evol. 2002; 54: 156–165.

16. Nagy PD, Simon AE. New insights into the mechanisms of RNA recombination. Virology. 1997; 235: 1–9.

17. Rowe CL, Fleming JO, Nathan MJ, Sgro JY, Palmenberg AC, Baker SC. Generation of coronavirus spike deletion variants by high-frequency recombination at regions of predicted RNA secondary structure. J. Virol. 1997; 71: 6183–90.

18. Cavanagh D. Coronaviridae: a review of coronaviruses and toroviruses. In: Schmidt A, Wolff MH, Weber O, editors. Coronaviruses with Special Emphasis on First Insights Concerning SARS. Birkhäuser-Verlag, 2005; pp. 1–54.

19. Lai MMC. RNA recombination in animal and plant viruses. Microbiological Reviews. 1992; 56: 61–79.

20. Pasternak AO, Spaan WJM, Snijder EJ. Nidovirus transcription: How to make sense…? Journal of General Virology. 2006; 87: 1403–1421.

21. Drosten C, Günther S, Preiser W, van der Werf S, Brodt HR, Becker S, et al. Identification of a Novel Coronavirus in Patients with Severe Acute Respiratory Syndrome. N. Engl. J. Med. 2003; 348: 1967–1976.

22. Ksiazek TG, Erdman D, Goldsmith CS, Zaki SR, Peret T, Emery S, et al. A Novel Coronavirus Associated with Severe Acute Respiratory Syndrome. N. Engl. J. Med. 2003; 348: 1953–1966.

23. Zaki AM, van Boheemen S, Bestebroer TM, Osterhaus ADME, Fouchier RAM. Isolation of a Novel Coronavirus from a Man with Pneumonia in Saudi Arabia. N. Engl. J. Med. 2012; 367: 1814–1820.

24. Guan Y, Zheng BJ, He YQ, Liu XL, Zhuang ZX, Cheung CL, et al. Isolation and characterization of viruses related to the SARS coronavirus from animals in Southern China. Science. 2003; 302: 276–278.

25. Alagaili AN, Briese T, Mishra N, Kapoor V, Sameroff SC, de Wit E, et al. Middle east respiratory syndrome coronavirus infection in dromedary camels in Saudi Arabia. MBio. 2014; 5.

26. Zhu N, Zhang D, Wang W, Li X, Yang Bo, Song J, et al. A Novel Coronavirus from Patients with Pneumonia in China, 2019. N. Engl. J. Med. 2020; doi:10.1056/NEJMoa2001017.

27. Lu H, Stratton CW, Tang Y. Outbreak of Pneumonia of Unknown Etiology in Wuhan China: the Mystery and the Miracle. J. Med. Virol. 2020; doi:10.1002/jmv.25678.

28. Hui DS, I Azhar E, Madani TA, Ntoumi F, Kock R, Dar O, et al. The continuing 2019-nCoV epidemic threat of novel coronaviruses to global health — The latest 2019 novel coronavirus outbreak in Wuhan, China. International Journal of Infectious Diseases. 2020; 91: 264–266.

29. Liu T, Hu J, Kang M, Lin L, Zhong H, Xiao J, et al. Transmission dynamics of 2019 novel coronavirus (2019-nCoV). BioRxiv [Preprint]. 2020 bioRxiv 919787 [posted 2020 January 25; cited 2020 January 31]. Available from: https://www.biorxiv.org/content/10.1101/2020.01.25.919787v1 doi: 10.1101/2020.01.25.919787.

30. Perlman S. Another Decade, Another Coronavirus. N. Engl. J. Med. 2020; doi: 10.1056/NEJMe2001126.

31. Gralinski LE, Menachery VD. Return of the Coronavirus: 2019-nCoV. Viruses. 2020; 12: 135.

32. 2019-nCoV Global Cases by Johns Hopkins CSSE. 2020 February 6 [cited 6 February 2020]. In: JHU CSSE website [Internet]. Available from: https://gisanddata.maps.arcgis.com/apps/opsdashboard/index.html#/bda7594740fd40299423467b48e9ecf6.

33. Novel Coronavirus(2019-nCoV) Situation Report - 13. 2002 February 02 [cited 02 February 2020]. In: WHO website [Internet]. Available from: https://www.who.int/docs/default-source/coronaviruse/situation-reports/20200202-sitrep-13-ncov-v3.pdf.

34. Chan JFW, Yuan S, Kok KH, To KKW, Chu H, Yang J, et al. A familial cluster of pneumonia associated with the 2019 novel coronavirus indicating person-to-person transmission: a study of a family cluster. Lancet. 2020; doi:10.1016/S0140-6736(20)30154-9.

35. Hu B, Zeng LP, Yang XL, Ge XY, Zhang W, Li B, et al. Discovery of a rich gene pool of bat SARS-related coronaviruses provides new insights into the origin of SARS coronavirus. PLoS Pathog. 2017; 13.

36. Dong N, Yang X, Ye L, Chen K, Chan EWC, Yang M, Chen S. Genomic and protein structure modelling analysis depicts the origin and infectivity of 2019-nCoV, a new coronavirus which caused a pneumonia outbreak in Wuhan, China. BioRxiv [Preprint]. 2020 bioRxiv 913368 [posted 2020 January 22; cited 2020 January 31]. Available from: https://www.biorxiv.org/content/10.1101/2020.01.20.913368v2 doi:10.1101/2020.01.20.913368.

37. Guo Q, Li M, Wang C, Wang P, Fang Z, Tan J, et al. Host and infectivity prediction of Wuhan 2019 novel coronavirus using deep learning algorithm. BioRxiv [Preprint]. 2020 bioRxiv 914044 [posted 2020 January 22; cited 2020 January 31]. Available from: https://www.biorxiv.org/content/10.1101/2020.01.21.914044v2 doi: 10.1101/2020.01.21.914044.

38. Wu F, Zhao S, Yu B, Chen YM, Wang W, Hu Y, et al. Complete genome characterisation of a novel coronavirus associated with severe human respiratory disease in Wuhan, China. BioRxiv [Preprint]. 2020 bioRxiv 919183 [posted 2020 February 02; cited 2020 February 02]. Available from: https://www.biorxiv.org/content/10.1101/2020.01.24.919183v2 doi:10.1101/2020.01.24.919183.

39. Paraskevis D, Kostaki EG, Magiorkinis G, Panayiotakopoulos G, Tsiodras S. Full-genome evolutionary analysis of the novel corona virus (2019-nCoV) rejects the hypothesis of emergence as a result of a recent recombination event. BioRxiv [Preprint]. 2020 bioRxiv 920249 [posted 2020 January 27; cited 2020 January 31]. Available from: https://www.biorxiv.org/content/10.1101/2020.01.26.920249v1 doi: 10.1101/2020.01.26.920249.

40. Ji W, Wang W, Zhao X, Zai J, Li X. Homologous recombination within the spike glycoprotein of the newly identified coronavirus may boost cross species transmission from snake to human. J. Med. Virol. 2020; doi:10.1002/jmv.25682.

41. Zhou P, Yang XL, Wang XG, Hu B, Zhang L, Zhang W, et al. Discovery of a novel coronavirus associated with the recent pneumonia outbreak in humans and its potential bat origin. BioRxiv [Preprint]. 2020 bioRxiv 914952 [posted 2020 January 23; cited 2020 January 31]. Available from: https://www.biorxiv.org/content/10.1101/2020.01.22.914952v1 doi:10.1101/2020.01.22.914952.

42. Letko M, Munster V. Functional assessment of cell entry and receptor usage for lineage B *β*-coronaviruses, including 2019-nCoV. BioRxiv [Preprint]. 2020 bioRxiv 915660 [posted 2020 January 22; cited 2020 January 31]. Available from: https://www.biorxiv.org/content/10.1101/2020.01.22.915660v1 doi:10.1101/2020.01.22.915660.

43. Zhao Y, Zhao Z, Wang Y, Zhou Y, Ma Y, Zuo W. Single-cell RNA expression profiling of ACE2, the putative receptor of Wuhan 2019-nCoV. BioRxiv [Preprint]. 2020 bioRxiv 919985 [posted 2020 January 26; cited 2020 January 31]. Available from: https://www.biorxiv.org/content/10.1101/2020.01.26.919985v1 doi:10.1101/2020.01.26.919985.

44. Li Y, Zhang J, Wang N, Li H, Shi Y, Gui G, et al. Therapeutic Drugs Targeting 2019-nCoV Main Protease by High-Throughput Screening. BioRxiv [Preprint]. 2020 bioRxiv 922922 [posted 2020 January 30; cited 2020 January 31]. Available from: https://www.biorxiv.org/content/10.1101/2020.01.28.922922v2 doi:10.1101/2020.01.28.922922.

45. Liu X, Wang XJ. Potential inhibitors for 2019-nCoV coronavirus M protease from clinically approved medicines. BioRxiv [Preprint]. 2020 bioRxiv 924100 [posted 2020 January 29; cited 2020 January 31]. Available from: https://www.biorxiv.org/content/10.1101/2020.01.29.924100v1 doi:10.1101/2020.01.29.924100.

46. Vinga S, Almeida J. Alignment-free sequence comparison-a review. Bioinformatics. 2003; 19(4): 513–523.

47. Zielezinski A, Vinga S, Almeida J, Karlowski WM. Alignment-free sequence comparison: benefits, applications, and tools. Genome Biology. 2017, 18: 186.

48. Kari L, Hill KA, Sayem AS, Karamichalis R, Bryans N, Davis K, Dattani NS. Mapping the space of genomic signatures. PLoS ONE. 2015; 10: e0119815.

49. Karamichalis R, Kari L, Konstantinidis S, Kopecki S. An investigation into inter- and intragenomic variations of graphic genomic signatures. BMC Bioinformatics. 2015; 16: 246.

50. Solis-Reyes S, Avino M, Poon A. An open-source k-mer based machine learning tool for fast and accurate subtyping of HIV-1 genomes. PLoS ONE. 2018; 13: e0206409.

51. Randhawa GS, Hill KH, Kari L. ML-DSP: Machine Learning with Digital Signal Processing for ultrafast, accurate, and scalable genome classification at all taxonomic levels. BMC Genomics. 2019; 20: 267.

52. Randhawa GS, Hill KH, Kari L. MLDSP-GUI: an alignment-free standalone tool with an interactive graphical user interface for DNA sequence comparison and analysis. Bioinformatics. 2019; btz918.

53. Jeffre HJ. Chaos game representation of gene structure. Nucleic Acids Res. 1990; 18: 2163–2170.

54. Asuero AG, Sayago A, González AG. The correlation coefficient: an overview. Crit Rev Anal Chem. 2006; 36(1): 41–59.

55. Karamichalis R, Kari L. MoDMaps3D: an interactive webtool for the quantification and 3D visualization of interrelationships in a dataset of DNA sequences. Bioinformatics. 2017; 33(19): 3091–3.

56. Kruskal J. Multidimensional scaling by optimizing goodness of fit to a nonmetric hypothesis. Psychometrika. 1964; 29: 1–27.

57. Carneiro RL, Requião RD, Rossetto S, Domitrovic T, Palhano FL. Codon stabilization coefficient as a metric to gain insights into mRNA stability and codon bias and their relationships with translation. Nucleic acids research. 2019; 47(5): 2216–2228.

58. Karumathil S, Raveendran NT, Ganesh D, Kumar NS, Nair RR, Dirisala VR. Evolution of Synonymous Codon Usage Bias in West African and Central African Strains of Monkeypox Virus. Evolutionary Bioinformatics Online. 2018; 14: doi:10.1177/1176934318761368.

59. Vinogradov AE, Anatskaya OV. DNA helix: the importance of being AT-rich. Mammalian Genome. 2017; 9(10): 455–464.

60. Hollander M, Wolfe DA, Chicken E. Nonparametric statistical methods, 3rd Edition, John Wiley & Sons; 2013.

61. Zhao S, Lin Q, Ran J, Musa SS, Yang G, Wang W, et al. Preliminary estimation of the basic reproduction number of novel coronavirus (2019-nCoV) in China, from 2019 to 2020: A data-driven analysis in the early phase of the outbreak. International Journal of Infectious Diseases [In Press] [received 2020 January 23; revised 2020 January 27; accepted 2020 January 27; cited 2020 February 1], 2020.

62. Shao P, Shan Y. Beware of asymptomatic transmission: Study on 2019-nCoV prevention and control measures based on extended SEIR model. BioRxiv [Preprint]. 2020 bioRxiv 923169 [posted 2020 January 28; cited 2020 January 31]. Available from: https://www.biorxiv.org/content/10.1101/2020.01.28.923169v1 doi:10.1101/2020.01.28.923169.

63. Chen Z, Zhang W, Lu Y, Guo C, Guo Z, Liao C, et al. From SARS-CoV to Wuhan 2019-nCoV Outbreak: Similarity of Early Epidemic and Prediction of Future Trends. BioRxiv [Preprint]. 2020 bioRxiv 919241 [posted 2020 January 27; cited 2020 January 31]. Available from: https://www.biorxiv.org/content/10.1101/2020.01.24.919241v3 doi:10.1101/2020.01.24.919241.

64. Hayward JA, Tachedjian M, Cui J, Field H, Holmes EC, Wang L, Tachedjian G. Identification of diverse full-length endogenous betaretroviruses in megabats and microbats. Retrovirology. 2013; 10.

65. Cui J, Tachedjian G, Wang LF. Bats and Rodents Shape Mammalian Retroviral Phylogeny. Sci. Rep. 2015; 5.

66. Hayward JA, Tachedjian M, Cui J, Cheng AZ, Johnson A, Baker ML, et al. Differential evolution of antiretroviral restriction factors in pteropid bats as revealed by APOBEC3 gene complexity. Mol. Biol. Evol. 2018; 35: 1626–1637.

67. Wong A, Li X, Lau S, Woo P. Global Epidemiology of Bat Coronaviruses. Viruses. 2019; 11(2): 174.

68. Yang XL, Hu B, Wang B, Wang MN, Zhang Q, Zhang W, et al. Isolation and Characterization of a Novel Bat Coronavirus Closely Related to the Direct Progenitor of Severe Acute Respiratory Syndrome Coronavirus. J. Virol. 2016; 90: 3253–3256.

69. Lau SK, Li KS, Tsang AK, Lam CS, Ahmed S, Chen H, et al. Genetic Characterization of Betacoronavirus Lineage C Viruses in Bats Reveals Marked Sequence Divergence in the Spike Protein of Pipistrellus Bat Coronavirus HKU5 in Japanese Pipistrelle: Implications for the Origin of the Novel Middle East Respiratory Syndrome Coronavirus. J. Virol. 2013; 87: 8638–8650.

70. Lacroix A, Duong V, Hul V, San S, Davun H, Omaliss K, et al. Genetic diversity of coronaviruses in bats in Lao PDR and Cambodia. Infect. Genet. Evol. 2017; 48: 10–18.

71. Drexler JF, Gloza-Rausch F, Glende J, Corman VM, Muth D, Goettsche M, et al. Genomic Characterization of Severe Acute Respiratory Syndrome-Related Coronavirus in European Bats and Classification of Coronaviruses Based on Partial RNA-Dependent RNA Polymerase Gene Sequences. J. Virol. 2010; 84: 11336–11349.

72. Rihtarič D, Hostnik P, Steyer A, Grom J, Toplak I. Identification of SARS-like coronaviruses in horseshoe bats (Rhinolophus hipposideros) in Slovenia. Arch. Virol. 2010; 155: 507–514.

73. He B, Zhang Y, Xu L, Yang W, Yang F, Yun Feng, et al. Identification of Diverse Alphacoronaviruses and Genomic Characterization of a Novel Severe Acute Respiratory Syndrome-Like Coronavirus from Bats in China. J. Virol. 2014; 88: 7070–7082.

74. Wacharapluesadee S, Duengkae P, Rodpan A, Kaewpom T, Maneeorn P, Kanchanasaka B, et al. Diversity of coronavirus in bats from Eastern Thailand Emerging viruses. Virol. J. 2015; 12: 1–7.

75. Tong S, Conrardy C, Ruone S, Kuzmin IV, Guo X, Tao Y, et al. Detection of novel SARS-like and other coronaviruses in bats from Kenya. Emerg. Infect. Dis. 2009; 15: 482–485.

76. Lau SKP, Woo PCY, Li KSM, Huang Y, Tsoi H, Wong BHL, et al. Severe acute respiratory syndrome coronavirus-like virus in Chinese horseshoe bats. Proc. Natl. Acad. Sci. 2005; National Academy of Sciences, U. S. A., 102: 14040–14045.

77. Virologists weigh in on novel coronavirus in China’s outbreak. 2020 January 08 [cited 31 January 2020]. In: University of Minnesota [Internet]. Available from: http://www.cidrap.umn.edu/news-perspective/2020/01/virologists-weigh-novel-coronavirus-chinas-outbreak.

78. nCoV’s relationship to bat coronaviruses & recombination signals (no snakes) - no evidence the 2019-nCoV lineage is recombinant. 2020 January 31 [cited 31 January 2020]. In: Virological blog [Internet]. Available from: http://virological.org/t/ncovs-relationship-to-bat-coronaviruses-recombination-signals-no-snakes-no-evidence-the-2019-nCoV-lineage-is-recombinant/331.

79. Experts: nCoV spread in China’s cities could trigger global epidemic. 2020 January 27 [cited 31 January 2020]. In: University of Minnesota [Internet]. Available from: http://www.cidrap.umn.edu/news-perspective/2020/01/experts-ncov-spread-chinas-cities-could-trigger-global-epidemic.

80. China detects large quantity of novel coronavirus at Wuhan seafood market. 2020 January 27 [cited 31 January 2020]. In: Xinhuanet News [Internet]. Available from: http://www.xinhuanet.com/english/2020-01/27/c_138735677.htm.

81. Takata MA, Gonçalves-Carneiro D, Zang TM, Soll SJ, York A, Blanco-Melo D, Bieniasz PD. CG dinucleotide suppression enables antiviral defence targeting non-self RNA. Nature. 2017; 550(7674): 124–127.

82. Greenbaum BD, Levine AJ, Bhanot G, Rabadan R. Patterns of evolution and host gene mimicry in influenza and other RNA viruses. PLoS Pathogens. 2008; 4(6): doi:10.1371/journal.ppat.1000079.

83. Lobo FP, Mota BEF, Pena SDJ, Azevedo V, Macedo AM, Tauch A, et al. Virus-host coevolution: Common patterns of nucleotide motif usage in Flaviviridae and their hosts. PLoS ONE. 2009; 4(7): 10.1371/journal.pone.0006282.

84. Kindler E, Thiel V. To sense or not to sense viral RNA-essentials of coronavirus innate immune evasion. Current Opinion in Microbiology. 2014; 20: 68–75.

85. Milewska A, Kindler E, Vkovski P, Zeglen S, Ochman M, Thiel V, et al. APOBEC3-mediated restriction of RNA virus replication. Scientific Reports. 2018; 8(1): doi:10.1038/s41598-018-24448-2.

86. Bishop KN, Holmes RK, Sheehy AM, Malim MH. APOBEC-mediated editing of viral RNA. Science. 2004; 305(5684): 645.

87. Pyrc K, Jebbink MF, Berkhout B, Van der Hoek L. Genome structure and transcriptional regulation of human coronavirus NL63. Virology Journal. 2004; 1(1): 7.

88. Berkhout B, Van Hemert F. On the biased nucleotide composition of the human coronavirus RNA genome. Virus Research. 2015; 202: 41–47.

89. Woo PCY, Lau SKP, Huang Y, Yuen KY. Coronavirus diversity, phylogeny and interspecies jumping. Experimental Biology and Medicine. 2009; 234(10): 1117–1127.

90. Woo PCY, Huang Y, Lau SKP, Yuen KY. Coronavirus Genomics and Bioinformatics Analysis. Viruses. 2010; 2(8): 1804–1820.

91. Xue X, Yu H, Yang H, Xue F, Wu Z, Shen W, et al. Structures of Two Coronavirus Main Proteases: Implications for Substrate Binding and Antiviral Drug Design. J. Virol. 2008; 82: 2515–2527.

92. Anand K, Ziebuhr J, Wadhwani P, Mesters JR, Hilgenfeld R. Coronavirus main proteinase (3CLpro) Structure: Basis for design of anti-SARS drugs. Science. 2003; 300: 1763–1767.

93. Nukoolkarn V, Lee VS, Malaisree M, Aruksakulwong O, Hannongbua S. Molecular dynamic simulations analysis of ritronavir and lopinavir as SARS-CoV 3CLpro inhibitors. J. Theor. Biol. 2008; 254: 861–867.

94. Xu Z, Peng C, Shi Y, Zhu Z, Mu K, Wang X, Zhu W. Nelfinavir was predicted to be a potential inhibitor of 2019-nCoV main protease by an integrative approach combining homology modelling, molecular docking and binding free energy calculation. BioRxiv [Preprint]. 2020 bioRxiv 921627 [posted 2020 January 28; cited 2020 January 31]. Available from: https://www.biorxiv.org/content/10.1101/2020.01.27.921627v1 doi: 10.1101/2020.01.27.921627.

