## Supplementary material for "Machine learning using intrinsic genomic signatures for rapid classification of novel pathogens: COVID-19 case study": S1 Table.; S2 Table.; S3 Table.

**Gurjit S. Randhawa, Maximillian P.M. Soltysiak, Hadi El Roz, Camila P.E. de Souza, Kathleen A. Hill, and Lila Kari**

### 1) Software availability

MLDSP-GUI is an open-source alignment-free tool with Graphical User Interface is publicly available for download at the following link (No license is required to download and use the tool):

https://sourceforge.net/projects/mldsp-gui/

MLDSP is an open-source alignment-free tool (MATLAB license required to run this program) available at the following link:

https://github.com/grandhawa/MLDSP

### 2) Spearman's rank correlation coefficient test results

The ρ values from the Spearman's correlation coefficient test for k=1 to k=7 are given in Supplementary Table S1. The P-value is <1e-5 for k=2 to k=6, 0.0833 for k=1 with an exception of 0.3333 in case of *Deltacorovavirus*.

| 2019-nCoV vs. | k=1 | k=2 | k=3 | k=4 | k=5 | k=6 | k=7 |
|---|---|---|---|---|---|---|---|
| ***Alphacoronavirus*** | 1 | 0.97 | 0.96 | 0.96 | 0.94 | 0.88 | 0.7 |
| ***Betacoronavirus*** | 1 | 0.95 | 0.95 | 0.95 | 0.94 | 0.89 | 0.74 |
| ***Gammacoronavirus*** | 1 | 0.93 | 0.91 | 0.92 | 0.90 | 0.83 | 0.63 |
| ***Deltacoronavirus*** | 0.8 | 0.98 | 0.94 | 0.92 | 0.90 | 0.81 | 0.60 |
| ***Embecovirus*** | 1 | 0.85 | 0.88 | 0.88 | 0.86 | 0.79 | 0.59 |
| ***Merbecovirus*** | 1 | 0.96 | 0.95 | 0.94 | 0.92 | 0.84 | 0.64 |
| ***Nobecovirus*** | 1 | 0.89 | 0.83 | 0.83 | 0.8 | 0.73 | 0.54 |
| ***Sarbecovirus*** | 1 | 0.98 | 0.97 | 0.97 | 0.95 | 0.88 | 0.72 |

**Supplementary Table S1:** Spearman's rank correlation coefficient (ρ) value for k=1 to k=7.

#### 3) Dataset availability

The sequences are downloaded from three databases: Virus-Host-DB, NCBI, and GISAID.

***Virus-Host-DB:***
All the viral sequences (apart from the ones downloaded from NCBI and GISAID) used in this study are obtained from the Virus-Host-DB available at:
https://www.genome.jp/virushostdb/

***NCBI***
Wuhan-Hu-1 complete genome (Accession: NC_045512.2), bat-SL-CoVZC45 (Accession: MG772933.1), and bat-SL-CoVZXC21 (Accession: MG772934.1) are obtained from the Virus-Host-DB available at:
https://www.ncbi.nlm.nih.gov/

***GISAID***
28 2019-nCoV sequences and Beta-CoV-RaTG13 sequence (from Bat) are downloaded from the GISAID. We gratefully acknowledge the Authors, the Originating and Submitting Laboratories for their sequence and metadata shared through GISAID, which is used in this research. All submitters of 28 2019-nCoV sequences and 1 BetaCoV/Bat/RaTG13 sequence may be contacted directly via www.gisaid.org

| Accession ID | Originating lab | Submitting lab | Authors |
|---|---|---|---|
| EPI_ISL_404227 | Zhejiang Provincial Center for Disease Control and Prevention | Department of Microbiology, Zhejiang Provincial Center for Disease Control and Prevention | Yin Chen, Yanjun Zhang, Haiyan Mao, Junhang Pan, Xiuyu Lou, Yiyu Lu, Juying Yan, Hanping Zhu, Jian Gao, Yan Feng, Yi Sun, Hao Yan, Zhen Li, Yisheng Sun, Liming Gong, Qiong Ge, Wen Shi, Xinying Wang, Wenwu Yao, Zhangnv Yang, Fang Xu, Chen Chen, Enfu Chen, Zhen Wang, Zhiping Chen, Jianmin Jiang, Chonggao Hu |
| EPI_ISL_404228 | Zhejiang Provincial Center for Disease Control and Prevention | Department of Microbiology, Zhejiang Provincial Center for Disease Control and Prevention | Yanjun Zhang, Yin Chen, Haiyan Mao, Junhang Pan, Xiuyu Lou, Yiyu Lu, Juying Yan, Hanping Zhu, Jian Gao, Yan Feng, Yi Sun, Hao Yan, Zhen Li, Yisheng Sun, Liming Gong, Qiong Ge, Wen Shi, Xinying Wang, Wenwu Yao, Zhangnv Yang, Fang Xu, Chen Chen, Enfu Chen, Zhen Wang, Zhiping Chen, Jianmin Jiang, Chonggao Hu |
| EPI_ISL_402132 | Wuhan Jinyintan Hospital | Hubei Provincial Center for Disease Control and Prevention | Bin Fang, Xiang Li, Xiao Yu, Linlin Liu, Bo Yang, Faxian Zhan, Guojun Ye, Xixiang Huo, Junqiang Xu, Bo Yu, Kun Cai, Jing Li, Yongzhong Jiang. |
| EPI_ISL_402127<br>EPI_ISL_402128<br>EPI_ISL_402129<br>EPI_ISL_402130<br>EPI_ISL_402124 | Wuhan Jinyintan Hospital | Wuhan Institute of Virology, Chinese Academy of Sciences | Peng Zhou, Xing-Lou Yang, Ding-Yu Zhang, Lei Zhang, Yan Zhu, Hao-Rui Si, Zhengli Shi |
| EPI_ISL_403963<br>EPI_ISL_403962 | Bamrasnaradura Hospital | 1. Department of Medical Sciences, Ministry of Public Health, Thailand 2. Thai Red Cross | Pilailuk, Okada; Siripaporn, Phuygun; Thanutsapa, Thanadachakul; Supaporn, Wacharapluesadee; Sittiporn,Parnmen; Warawan,Wongboot; Sunthareeya, |

| | | | |
|---|---|---|---|
| | | Emerging Infectious Diseases - Health Science Centre 3. Department of Disease Control, Ministry of Public Health, Thailand | Waicharoen; Rome, Buathong; Malinee, Chittaganpitch; Nanthawan, Mekha |
| EPI_ISL_402120<br>EPI_ISL_402119<br>EPI_ISL_402121 | National Institute for Viral Disease Control and Prevention, China CDC | National Institute for Viral Disease Control and Prevention, China CDC | Wenjie Tan, Xiang Zhao, Wenling Wang, Xuejun Ma, Yongzhong Jiang , Roujian Lu, Ji Wang, Weimin Zhou , Peihua Niu, Peipei Liu, Faxian Zhan, Weifeng Shi, Baoying Huang , Jun Liu, Li Zhao, Yao Meng, Xiaozhou He, Fei Ye, Na Zhu, Yang Li, Jing Chen, Wenbo Xu, George F. Gao, Guizhen Wu |
| EPI_ISL_402123 | Institute of Pathogen Biology, Chinese Academy of Medical Sciences & Peking Union Medical College | Institute of Pathogen Biology, Chinese Academy of Medical Sciences & Peking Union Medical College | Lili Ren, Jianwei Wang, Qi Jin, Zichun Xiang, Zhiqiang Wu, Chao Wu, Yiwei Liu |
| EPI_ISL_402125 | unknown | National Institute for Communicable Disease Control and Prevention (ICDC) Chinese Center for Disease Control and Prevention (China CDC) | Zhang,Y.-Z., Wu,F., Chen,Y.-M., Pei,Y.-Y., Xu,L., Wang,W., Zhao,S., Yu,B., Hu,Y., Tao,Z.-W., Song,Z.-G., Tian,J.-H., Zhang,Y.-L., Liu,Y., Zheng,J.-J., Dai,F.-H., Wang,Q.-M., She,J.-L. and Zhu,T.-Y. |
| EPI_ISL_403931<br>EPI_ISL_403928<br>EPI_ISL_403930<br>EPI_ISL_403929 | Institute of Pathogen Biology, Chinese Academy of Medical Sciences & Peking Union Medical College | Institute of Pathogen Biology, Chinese Academy of Medical Sciences & Peking Union Medical College | Lili Ren, Jianwei Wang, Qi Jin, Zichun Xiang, Zhiqiang Wu, Chao Wu, Yiwei Liu |
| EPI_ISL_403937<br>EPI_ISL_403936<br>EPI_ISL_403935<br>EPI_ISL_403934<br>EPI_ISL_403933<br>EPI_ISL_403932 | Guangdong Provincial Center for Diseases Control and Prevention; Guangdong Provincial Public Health | Department of Microbiology, Guangdong Provincial Center for Diseases Control and Prevention | Min Kang, Jie Wu, Jing Lu, Tao Liu, Baisheng Li, Shujiang Mei, Feng Ruan, Lifeng Lin, Changwen Ke, Haojie Zhong, Yingtao Zhang, Lirong Zou, Xuguang Chen, Qi Zhu, Jianpeng Xiao, Jianxiang Geng, Zhe Liu, Jianxiong Hu, Weilin Zeng, Xing Li, Yuhuang Liao, Xiujuan Tang, Songjian Xiao, Ying Wang, Yingchao Song, Xue Zhuang, Lijun Liang, Guanhao He, Huihong Deng, Tie Song, Jianfeng He, Wenjun Ma |
| EPI_ISL_404895 | Providence Regional Medical Center | Division of Viral Diseases, Centers for Disease Control and Prevention | Queen,K., Tao,Y., Li,Y., Paden,C.R., Lu,X., Zhang,J., Gerber,S.I., Lindstrom,S. |

| | | | |
|---|---|---|---|
| EPI_ISL_404253 | IL Department of Public Health Chicago Laboratory | Pathogen Discovery, Respiratory Viruses Branch, Division of Viral Diseases, Centers for Dieases Control and Prevention | Ying Tao, Krista Queen, Clinton R. Paden, Jing Zhang, Yan Li, Anna Uehara, Xiaoyan Lu, Brian Lynch, Senthil Kumar K. Sakthivel, Brett L. Whitaker, Shifaq Kamili, Lijuan Wang, Janna' R. Murray, Susan I. Gerber, Stephen Lindstrom, Suxiang Tong |
| EPI_ISL_405839 | The University of Hong Kong - Shenzhen Hospital | Li Ka Shing Faculty of Medicine, The University of Hong Kong | Chan,J.F.-W., Yuan,S., Kok,K.H., To,K.K.-W., Chu,H., Yang,J., Xing,F., Liu,J., Yip,C.C.-Y., Poon,R.W.-S., Tsai,H.W., Lo,S.K.-F., Chan,K.H., Poon,V.K.-M., Chan,W.M., Ip,J.D., Cai,J.P., Cheng,V.C.-C., Chen,H., Hui,C.K.-M. and Yuen,K.Y. |
| EPI_ISL_402131 (Bat RaTG13) | Wuhan Institute of Virology, Chinese Academy of Sciences | Wuhan Institute of Virology, Chinese Academy of Sciences | Yan Zhu, Ping Yu, Bei Li, Ben Hu, Hao-Rui Si, Xing-Lou Yang, Peng Zhou, Zheng-Li Shi |

**Supplementary Table S2:** Accession IDs of the Sequences downloaded from the GISAID.

The accession numbers and the sources for all the used sequences are given below:

| **Test-1; Source: Virus-Host-DB** |
|---|
| **Adenoviridae** |
| AB448767,AC_000007,JN880453,JN880454,JN880455,JN880456,JN935766,JQ326209,JQ776547,KC529648,KC693021,KF268207,AC_000008,KF279629,KF528688,KF802426,KF906413,KM591901,KM591902,KM591903,NC_000899,NC_000942,NC_001405,AC_000009,NC_001454,NC_001460,NC_001720,NC_001734,NC_001813,NC_001876,NC_001958,NC_002501,NC_002513,NC_002685,AC_000010,NC_002702,NC_003266,NC_004037,NC_006144,NC_006879,NC_009989,NC_010956,NC_011202,NC_011203,NC_012584,AC_000011,NC_012959,NC_014564,NC_014899,NC_014969,NC_015225,NC_015323,NC_015455,NC_015932,NC_016437,NC_016895,AC_000012,NC_017825,NC_017979,NC_020074,NC_020485,NC_020487,NC_021168,NC_021221,NC_022266,NC_022612,NC_022613,AC_000013,NC_024150,NC_024474,NC_024486,NC_024684,NC_025678,NC_025962,NC_027705,NC_027708,NC_028103,NC_028105,AC_000014,NC_028107,NC_028113,NC_029898,NC_029899,NC_029902,NC_030116,NC_030792,NC_030860,NC_030874,NC_031503,AC_000016,NC_031948,NC_032105,NC_034382,NC_034626,NC_034834,NC_035072,NC_035207,NC_035619,NC_038332,NC_038333,AC_000017,NC_038334,NC_039032,NC_040811,NC_043094,NC_043405,NC_043696,U46933,X73487,Y09598,AB724351,AC_000018,AC_000019,AC_000020,AC_000189,AC_000190,AC_000191,AF036092,AF083975,AF108105,AM749299,AB765926,AP012285,AP012302,AY458656,AY737797,AY737798,AY803294,AY849321,AY875648,DQ086466,DQ315364,AC_000001,DQ393829,DQ792570,DQ900900,DQ923122,EF121005,EF564601,FJ025899,FJ025900,FJ025901,FJ025902,AC_000002,FJ025903,FJ025904,FJ025905,FJ025906,FJ025907,FJ025908,FJ025909,FJ025910,FJ025911,FJ025912,AC_000003,FJ025913,FJ025914,FJ025915,FJ025916,FJ025917,FJ025918,FJ025919,FJ025920,FJ025921,FJ025922,AC_000004,FJ025923,FJ025924,FJ025925,FJ025926,FJ025927,FJ025928,FJ025929,FJ025930,FJ349096,FJ404771,AC_000005,FJ597732,FJ643676,FJ824826,GQ384080,GU191019,HM770721,HQ241818,HQ241820,HQ883276,JF964962,AC_000006,JN860676,JN860677,JN860678,JN860679,JN860680,JN880448,JN880449,JN880450,JN880451,JN880452 |
| **Anelloviridae** |
| AM711976,AM712003,AM712004,AM712030,AM712031,AM712032,AM712033,AM712034,FR823283,GU450331,HQ335082,HQ335083,HQ335084,HQ335085,JN704611,KJ194622,KM262781,KM262785,NC_001427,NC_002076,NC_002195,NC_007013,NC_007014,NC_009225,NC_012126,NC_014068,NC_014069,NC_014070,NC_014071,NC_014072,NC_014073,NC_014074,NC_014075,NC_014076,NC_014077,NC_014078,NC_014079,NC_014080,NC_014081,NC_014082,NC_014083,NC_014084,NC_0140 |

| |
|---|
| 85,NC_014086,NC_014087,NC_014088,NC_014089,NC_014090,NC_014091,NC_014092,NC_014093,NC_014094,NC_014095,NC_014096,NC_014097,NC_014480,NC_015212,NC_015396,NC_015783,NC_017091,NC_018401,NC_020498,NC_022788,NC_022789,NC_024890,NC_024891,NC_024908,NC_025215,NC_025726,NC_025727,NC_025966,NC_026138,NC_026662,NC_026663,NC_026664,NC_026764,NC_026765,NC_027059,NC_027430,NC_030297,NC_030650,NC_034978,NC_035135,NC_035136,NC_035192,NC_038336,NC_038337,NC_038338,NC_038339,NC_038340,NC_038341,NC_038342,NC_038343,NC_038344,NC_038345,NC_038346,NC_038347,NC_038348,NC_038349,NC_038350,NC_038351,NC_038352,NC_038353,NC_038354,NC_038355,NC_038356,NC_038357,NC_038358,NC_038359,NC_038360,NC_038361,NC_038362,NC_038363,NC_040531,NC_040546,NC_040547,NC_040617,NC_040618,NC_040668,NC_040686,NC_040687,NC_040720,NC_040801,NC_043413,NC_043414,NC_043415 |
| **Caudovirales** |
| AB626963,AB746912,AB757801,AF527608,AP011956,AY526908,AY526909,CP000711,CP008753,DQ113772,DQ121662,DQ222851,DQ289556,DQ394806,DQ394807,DQ394808,DQ394809,DQ394810,DQ426905,DQ838728,EU056923,EU568876,EU622808,FQ482084,GQ303261,GQ478082,GQ478083,GQ478085,GQ478087,GU196281,HE614282,HE956707,HE983844,HG428758,HG793132,HG796219,HG796220,HG796221,HM152765,HQ110083,HQ634152,HQ641341,HQ641343,HQ641344,HQ641346,JF314845,JF767210,JF773396,JN175269,JN254801,JN255163,JN699002,JN811560,JQ067085,JQ267518,JQ691610,JQ740790,JQ740791,JQ740792,JQ740793,JQ740794,JQ740795,JQ740796,JQ740797,JQ740798,JQ740799,JQ740800,JQ740801,JQ740802,JQ740803,JQ740805,JQ740806,JQ740807,JQ740808,JQ740809,JQ740810,JQ740811,JQ740812,JQ740814,JQ780163,JQ957925,JQ965700,JQ965701,JQ965702,JQ965703,JX000007,JX174275,JX274646,JX274647,JX403939,JX409894,JX409895,JX421753,JX483873,JX483874,JX483875,JX483879,JX483880,JX564242,JX570703,JX570707,JX570708,JX570711,JX681814,KC182543,KC182544,KC182545,KC182548,KC182549,KC182550,KC330681,KC333879,KC348598,KC348599,KC348600,KC348601,KC348602,KC348603,KC348604,KC413987,KC413988,KC522412,KC542353,KC556893,KC556894,KC556895,KC556896,KC556898,KC787107,KC787108,KC821615,KC821627,KC911856,KC911857,KC969441,KF030445,KF302032,KF302033,KF302035,KF302036,KF302037,KF591601,KF669657,KF676640,KF751793,KF751794,KF751795,KF751796,KF751797,KF771236,KF800937,KJ018210,KJ021043,KJ417497,KJ502657,KJ545483,KJ572844,KJ578763,KJ578764,KJ578766,KJ578769,KJ578771,KJ578775,KJ578777,KJ617393,KJ725374,KM058087,KM091442,KM091443,KM091444,KM233455,KM591905,KM612260,KM612261,KM612262,KM612263,KM612265,KM923970,KP017310,KP209285,KP296794,KP791807,KP869108,KR131710,LK985321,LN610580,LN681534,M11813,NC_000871,NC_000872,NC_000896,NC_000929,NC_000935,NC_001271,NC_001317,NC_001416,NC_001604,NC_001609,NC_001629,NC_001697,NC_001706,NC_001825,NC_001835,NC_001895,NC_001900,NC_001901,NC_001902,NC_001909,NC_001978,NC_002072,NC_002166,NC_002167,NC_002185,NC_002214,NC_002321,NC_002371,NC_002486,NC_002515,NC_002519,NC_002628,NC_002649,NC_002661,NC_002666,NC_002667,NC_002668,NC_002669,NC_002670,NC_002671,NC_002703,NC_002730,NC_002747,NC_002796,NC_003050,NC_003085,NC_003157,NC_003216,NC_003278,NC_003288,NC_003291,NC_003298,NC_003313,NC_003315,NC_003356,NC_003390,NC_003444,NC_003524,NC_003907,NC_004066,NC_004112,NC_004165,NC_004166,NC_004167,NC_004302,NC_004303,NC_004305,NC_004313,NC_004333,NC_004348,NC_004456,NC_004466,NC_004584,NC_004585,NC_004586,NC_004587,NC_004588,NC_004589,NC_004615,NC_004616,NC_004617,NC_004664,NC_004665,NC_004678,NC_004679,NC_004740,NC_004745,NC_004746,NC_004775,NC_004777,NC_004814,NC_004821,NC_004827,NC_004831,NC_004902,NC_004996,NC_005045,NC_005056,NC_005069,NC_005178,NC_005263,NC_005294,NC_005340,NC_005342,NC_005344,NC_005345,NC_005354,NC_005355,NC_005356,NC_005357,NC_005822,NC_005833,NC_005841,NC_005879,NC_005882,NC_005884,NC_005886,NC_005887,NC_005891,NC_005893,NC_006356,NC_006548,NC_006557,NC_006882,NC_006936,NC_006940,NC_006949,NC_006953,NC_007019,NC_007046,NC_007047,NC_007048,NC_007049,NC_007050,NC_007051,NC_007052,NC_007053,NC_007054,NC_007055,NC_007056,NC_007057,NC_007058,NC_007059,NC_007060,NC_007061,NC_007062,NC_007063,NC_007064,NC_007065,NC_007145,NC_007149,NC_007291,NC_007456,NC_007458,NC_007497,NC_007501,NC_007603,NC_007637,NC_007709,NC_007710,NC_007734,NC_007804,NC_007805,NC_007806,NC_007807,NC_007808,NC_007814,NC_007924,NC_007967,NC_008152,NC_008193,NC_008201,NC_008202,NC_008265,NC_008363,NC_008364,NC_008367,NC_008370,NC_008371,NC_008376,NC_008583,NC_008617,NC_008689,NC_008694,NC_008695,NC_008717,NC_008721,NC_008722,NC_008723,NC_008798,NC_008799,NC_009014,NC_009016,NC_009018,NC_009232,NC_009234,NC_009235,NC_009236,NC_009237,NC_009382,NC_009514,NC_009526,NC_009531,NC_009540,NC_009541,NC_009542,NC_009543,NC_009551,NC_009552,NC_009554,NC_009603,NC_009604, |

| |
|---|
| NC_009643,NC_009737,NC_009761,NC_009762,NC_009763,NC_009799,NC_009810,NC_009812,NC_009813,NC_009814,NC_009815,NC_009818,NC_009819,NC_009875,NC_009935,NC_009936,NC_009990,NC_010147,NC_010179,NC_010275,NC_010325,NC_010326,NC_010342,NC_010353,NC_010363,NC_010463,NC_010495,NC_010807,NC_010808,NC_010945,NC_011038,NC_011040,NC_011042,NC_011043,NC_011045,NC_011046,NC_011048,NC_011085,NC_011104,NC_011107,NC_011142,NC_011201,NC_011216,NC_011222,NC_011267,NC_011291,NC_011308,NC_011318,NC_011344,NC_011373,NC_011534,NC_011551,NC_011589,NC_011611,NC_011612,NC_011613,NC_011614,NC_011645,NC_011646,NC_011801,NC_011802,NC_011976,NC_012223,NC_012418,NC_012419,NC_012662,NC_012742,NC_012753,NC_012756,NC_012784,NC_012788,NC_012884,NC_013055,NC_013059,NC_013152,NC_013153,NC_013154,NC_013155,NC_013195,NC_013594,NC_013597,NC_013598,NC_013599,NC_013600,NC_013638,NC_013643,NC_013644,NC_013645,NC_013646,NC_013647,NC_013648,NC_013649,NC_013651,NC_013696,NC_014229,NC_014460,NC_014900,NC_015158,NC_015159,NC_015208 |
| **Geminiviridae** |
| KF229718,KF229722,KF652077,KJ628309,KM189819,L14460,L14461,L39638,NC_000869,NC_000870,NC_000882,NC_001346,NC_001359,NC_001369,NC_001412,NC_001438,NC_001439,NC_001466,NC_001467,NC_001468,NC_001478,NC_001507,NC_001508,NC_001647,NC_001828,NC_001868,NC_001917,NC_001928,NC_001929,NC_001930,NC_001931,NC_001932,NC_001933,NC_001934,NC_001935,NC_001936,NC_001937,NC_001938,NC_001939,NC_001983,NC_001984,NC_002046,NC_002047,NC_002048,NC_002049,NC_002510,NC_002543,NC_002555,NC_002556,NC_002817,NC_002981,NC_002984,NC_002985,NC_003199,NC_003326,NC_003357,NC_003379,NC_003418,NC_003434,NC_003493,NC_003504,NC_003505,NC_003556,NC_003609,NC_003664,NC_003665,NC_003708,NC_003709,NC_003722,NC_003744,NC_003803,NC_003804,NC_003822,NC_003825,NC_003828,NC_003830,NC_003831,NC_003856,NC_003857,NC_003860,NC_003861,NC_003862,NC_003865,NC_003866,NC_003867,NC_003868,NC_003887,NC_003891,NC_003896,NC_003897,NC_003898,NC_004005,NC_004042,NC_004043,NC_004044,NC_004071,NC_004090,NC_004091,NC_004096,NC_004097,NC_004098,NC_004099,NC_004100,NC_004101,NC_004147,NC_004153,NC_004192,NC_004300,NC_004356,NC_004558,NC_004559,NC_004569,NC_004580,NC_004581,NC_004582,NC_004583,NC_004607,NC_004608,NC_004609,NC_004611,NC_004612,NC_004613,NC_004614,NC_004618,NC_004625,NC_004626,NC_004627,NC_004628,NC_004630,NC_004634,NC_004635,NC_004637,NC_004638,NC_004639,NC_004640,NC_004641,NC_004642,NC_004644,NC_004645,NC_004646,NC_004647,NC_004648,NC_004650,NC_004651,NC_004654,NC_004655,NC_004656,NC_004657,NC_004658,NC_004659,NC_004660,NC_004661,NC_004662,NC_004673,NC_004674,NC_004675,NC_004676,NC_004732,NC_004755,NC_004824,NC_004825,NC_005031,NC_005032,NC_005319,NC_005320,NC_005321,NC_005330,NC_005331,NC_005338,NC_005347,NC_005348,NC_005635,NC_005636,NC_005807,NC_005811,NC_005812,NC_005842,NC_005843,NC_005844,NC_005845,NC_005846,NC_005850,NC_005851,NC_005852,NC_005853,NC_005855,NC_006358,NC_006359,NC_006384,NC_006631,NC_006874,NC_006876,NC_006995,NC_007210,NC_007211,NC_007290,NC_007338,NC_007339,NC_007638,NC_007723,NC_007724,NC_007726,NC_007727,NC_007730,NC_007965,NC_007966,NC_008056,NC_008057,NC_008058,NC_008059,NC_008236,NC_008267,NC_008283,NC_008284,NC_008299,NC_008304,NC_008305,NC_008316,NC_008317,NC_008329,NC_008373,NC_008374,NC_008377,NC_008492,NC_008493,NC_008494,NC_008495,NC_008517,NC_008559,NC_008779,NC_008780,NC_008793,NC_008794,NC_009030,NC_009031,NC_009088,NC_009354,NC_009451,NC_009490,NC_009491,NC_009545,NC_009546,NC_009547,NC_009548,NC_009549,NC_009550,NC_009553,NC_009605,NC_009606,NC_009607,NC_009612,NC_009644,NC_009645,NC_009646,NC_009647,NC_010238,NC_010293,NC_010294,NC_010307,NC_010313,NC_010352,NC_010417,NC_010435,NC_010439,NC_010440,NC_010441,NC_010618,NC_010647,NC_010648,NC_010713,NC_010714,NC_010791,NC_010792,NC_010797,NC_010799,NC_010812,NC_010818,NC_010833,NC_010834,NC_010835,NC_010836,NC_010837,NC_010838,NC_010839,NC_010840,NC_010946,NC_010947,NC_010948,NC_010949,NC_010950,NC_010951,NC_010952,NC_010953,NC_011024,NC_011052,NC_011058,NC_011096,NC_011135,NC_011181,NC_011182,NC_011268,NC_011309,NC_011346,NC_011347,NC_011348,NC_011583,NC_011584,NC_011804,NC_011805,NC_011919,NC_012041,NC_012118,NC_012120,NC_012137,NC_012206,NC_012481,NC_012482,NC_012492,NC_012553,NC_012554,NC_012664,NC_012665,NC_012786,NC_012787,NC_013017,AB007990,AB110218,AB162141,AB192965,AB192966,AB236321,AB236323,AB236325,AB306314,AB439841,AB439842,AB921568,AF105975,AF112352,AF112353,AF141897,AF141922,AF173555,AF173556,AF241479,AF291705,AF291706,AF329886,AF329888,AF329889,AF379637,AF416741,AF416742,AF428255,AF490004,AF491306,AJ132574,AJ132575,AJ223505,AJ224504,AJ311031,AJ314739,AJ314740,AJ319674,AJ420316,AJ420317,AJ420318,AJ457823,AJ4 |

| |
|---|
| 57824,AJ457985,AJ457986,AJ496286,AJ496287,AJ512761,AJ512762,AJ543429,AJ564742,AJ564743,AJ566744,AJ579307,AJ579308,AJ781302,AJ810156,AJ810157,AJ849916,AJ865337,AJ971263,AM181683,AM183224,AM230634,AM230635,AM261326,AM421522,AM691745,AM712436,AM940137,AM980883,AM989927,AY036009,AY036010,AY090555,AY090556,AY090557,AY090558,AY190290,AY190291,AY650283,AY754814,D00200,D00201,D00940,DQ845787,DQ868525,EF011559,EF015778,EF190217,EF536859,EF536861,EF536868,EF536873,EF536876,EF536878,EF536886,EU024118,EU024119,EU024120,EU273816,EU273817,EU273818,EU365686,EU856366,FJ176701,FJ560719,FJ751234,FM179613,FM210034,FM877474,FN252890,FN256256,FN256257,FN256258,FN256259,FN256260,FN256261,FN256292,FN297834,FN401520,FN436001,GU001879,GU076440,GU076443,GU076445,GU076447,GU076449,GU076451,GU076452,GU076454,GU180085,GU256531,GU440580,GU732203,HE580236,HE616777,HE659516,HE659517,HE793429,HM140364,HM140365,HM140366,HM140368,HM140369,HM140370,HM140371,HM626516,HM626517,HM859902,HM859903,JF451352,JN676150,JN676151,JN680352,JN680353,JN989417,JN989425,JN989441,JN989446,JQ247188,JQ303121,JQ303122,JQ621843,JX082259,JX448368,JX911332,K02029,K02030,KC149941,KC172700,KC427995,KC476655,KF176552 |
| **Genomoviridae** |
| JN704610,KF371641,KF371642,KP133076,KP133077,KP133078,KP133079,KP133080,NC_013116,NC_023844,NC_023870,NC_023871,NC_023872,NC_024689,NC_024690,NC_024691,NC_024909,NC_025728,NC_025729,NC_025730,NC_025731,NC_025732,NC_025733,NC_025734,NC_025735,NC_025736,NC_025737,NC_025738,NC_025741,NC_026144,NC_026161,NC_026162,NC_026163,NC_026164,NC_026165,NC_026166,NC_026167,NC_026168,NC_026169,NC_026254,NC_026261,NC_026806,NC_026807,NC_026808,NC_026809,NC_026810,NC_026817,NC_026818,NC_027776,NC_027820,NC_027821,NC_028459,NC_028460,NC_030138,NC_030139,NC_030140,NC_030141,NC_030142,NC_030143,NC_030144,NC_030145,NC_030146,NC_030147,NC_030447,NC_030448,NC_030887,NC_033270,NC_033736,NC_033742,NC_033743,NC_033747,NC_035137,NC_035138,NC_035139,NC_035197,NC_035477,NC_037062,NC_038479,NC_038480,NC_038481,NC_038482,NC_038483,NC_038484,NC_038485,NC_038486,NC_038487,NC_038488,NC_038489,NC_038490,NC_038491,NC_038492,NC_038493,NC_038494,NC_038495,NC_038496,NC_038497,NC_038498,NC_038499,NC_038501,NC_038502,NC_040317,NC_040326,NC_040327,NC_040330,NC_040338,NC_040339,NC_040340,NC_040346,NC_040347,NC_040348,NC_040351,NC_040370,NC_040371,NC_040372,NC_040379 |
| **Microviridae** |
| AJ550635,AY751298,DQ079873,DQ079878,DQ079880,DQ079881,DQ079882,DQ079883,DQ079884,DQ079885,DQ079886,DQ079887,DQ079888,DQ079889,DQ079890,DQ079891,DQ079892,DQ079893,DQ079894,DQ079895,DQ079896,DQ079897,DQ079898,DQ079899,DQ079900,DQ079901,DQ079902,DQ079903,DQ079904,DQ079905,DQ079906,DQ079907,DQ079908,DQ079909,KC237308,KC821628,KC821631,KF044309,KF044310,KJ997912,M14428,NC_001330,NC_001420,NC_001422,NC_001730,NC_001741,NC_001998,NC_002180,NC_002194,NC_002643,NC_003438,NC_007461,NC_007817,NC_007818,NC_007819,NC_007820,NC_007821,NC_007822,NC_007823,NC_007824,NC_007825,NC_007827,NC_007856,NC_012868,NC_015785,NC_021797,NC_021805,NC_022790,NC_026013,NC_026665,NC_027633,NC_027634,NC_027635,NC_027636,NC_027637,NC_027638,NC_027639,NC_027640,NC_027641,NC_027642,NC_027643,NC_027644,NC_027645,NC_027646,NC_027647,NC_027648,NC_028993,NC_028994,NC_029012,NC_029014,NC_030458,NC_030472,NC_030476,NC_040328,NC_040329,NC_040341,NC_040342,NC_040349,NC_040350,NC_040373,NC_040374,NC_040375 |
| **Ortervirales** |
| AB187566,AB611707,AF033809,AF053745,AF126467,AF221065,AF411814,DQ093792,DQ241301,DQ241302,DQ365814,DQ399707,DQ451009,DQ822073,EF133960,EF494181,EU293537,EU523109,FJ195346,FN692043,HM210570,HQ154630,HQ246218,HQ540591,HQ540595,J01998,JF274252,JF908815,JN032736,JN134185,JQ303225,JX245014,KC802224,KF029431,KF313137,KJ668270,KJ668271,KP284572,M11841,M14008,M16605,M23385,NC_000858,NC_001343,NC_001362,NC_001364,NC_001402,NC_001403,NC_001407,NC_001408,NC_001413,NC_001414,NC_001436,NC_001450,NC_001452,NC_001463,NC_001482,NC_001488,NC_001494,NC_001497,NC_001499,NC_001500,NC_001501,NC_001502,NC_001503,NC_001506,NC_001511,NC_001514,NC_001549,NC_001550,NC_001574,NC_001618,NC_001634,NC_001648,NC_001654,NC_001702,NC_001722,NC_001724,NC_001725,NC_001739,NC_001802,NC_001815,NC_001831,NC_001839,NC_001866,NC_001867,NC_001885,NC_001914,NC_001940,NC_002201,NC_003031,NC_003059,NC_003138,NC_003323,NC_003378,NC_003381,NC_003382,NC_003498,NC_003554,NC_004036,NC_004324,NC_004450,NC_004455,NC_004540,NC_004994,NC_005947,NC_006934,NC_006955,NC_007002,NC_007003,NC_007015,NC_007654,NC_007 |

| |
|---|
| 815,NC_008017,NC_008018,NC_008034,NC_008094,NC_009010,NC_009424,NC_009889,NC_010737,NC_010738,NC_010820,NC_010955,NC_011097,NC_011546,NC_011592,NC_011800,NC_011920,NC_012728,NC_013262,NC_013455,NC_014474,NC_014648,NC_015116,NC_015228,NC_015328,NC_015502,NC_015503,NC_015504,NC_015505,NC_015506,NC_015507,NC_015655,NC_015784,NC_017830,NC_018105,NC_018505,NC_018616,NC_018858,NC_020999,NC_022365,NC_022517,NC_022518,NC_023153,NC_023485,NC_024301,NC_026020,NC_026238,NC_026472,NC_026819,NC_027117,NC_027131,NC_027924,NC_028462,NC_029303,NC_029852,NC_029853,NC_030205,NC_030462,NC_031326,NC_033738,NC_033739,NC_034252,NC_035472,NC_038378,NC_038379,NC_038380,NC_038381,NC_038382,NC_038512,NC_038669,NC_038858,NC_038922,NC_038923,NC_038986,NC_038987,NC_038995,NC_039022,NC_039023,NC_039024,NC_039025,NC_039026,NC_039027,NC_039028,NC_039029,NC_039030,NC_039031,NC_039085,NC_039228,NC_039238,NC_039242,NC_040461,NC_040462,NC_040552,NC_040622,NC_040635,NC_040692,NC_040693,NC_040712,NC_040807,NC_040808,NC_040809,NC_040841,NC_043194,NC_043195,NC_043382,NC_043445,NC_043491,NC_043523,NC_043534,NC_043535,U04327,U21247,U85505,U85506,U94692,X00255,X13744,X54482,X57540,Y07725,Y13051 |
| **Papillomaviridae** |
| AB027020,AB027021,AB211993,AB331650,AB331651,AB361563,AB543507,AB793779,AF020905,AF092932,AF151983,AF293960,AF349909,AJ400628,AJ620205,AJ620208,AJ620209,AJ620211,AY395706,AY904722,AY904723,AY904724,D21208,D90252,D90400,DQ080079,DQ080080,DQ080082,DQ080083,DQ090005,DQ098913,DQ098917,DQ180494,DQ344807,EF028290,EF117891,EF422221,EF558838,EF558839,EF558840,EF558841,EF558842,EF558843,EF591300,EU240895,EU360723,EU410347,EU410348,EU410349,EU490515,EU490516,EU493091,EU918769,FJ492742,FJ492743,FJ947080,FM955837,FM955838,FM955839,FM955840,FM955841,FM955842,FN547152,FN598907,FN677755,FN677756,FR751039,GQ180114,GQ227670,GQ244463,GQ246950,GQ246951,GQ845441,GQ845442,GQ845444,GQ845445,GQ845446,GU117620,GU117624,GU117629,GU117630,GU117633,GU129016,HE963025,HG939559,HM999990,HM999991,HM999993,HM999994,HM999995,HM999997,HM999998,HM999999,J04353,JF304766,JF304767,JF304768,JF800658,JF906559,JN171845,JN709469,JN709470,JN709471,JN709472,JQ798171,JX174438,JX899359,KC138720,KC470240,KC858265,KF006398,KF006400,KJ145795,KM085343,KM983393,KP276343,L41216,M12732,M12737,M14119,M20219,M32305,M62877,M73236,M74117,NC_001352,NC_001354,NC_001355,NC_001356,NC_001357,NC_001457,NC_001458,NC_001522,NC_001523,NC_001524,NC_001526,NC_001531,NC_001541,NC_001576,NC_001583,NC_001586,NC_001587,NC_001591,NC_001593,NC_001595,NC_001596,NC_001605,NC_001619,NC_001676,NC_001678,NC_001690,NC_001691,NC_001693,NC_001694,NC_001789,NC_002232,NC_003348,NC_003748,NC_003973,NC_004068,NC_004104,NC_004195,NC_004197,NC_004500,NC_004765,NC_005134,NC_006563,NC_006564,NC_006951,NC_007150,NC_007612,NC_008032,NC_008184,NC_008188,NC_008189,NC_008297,NC_008298,NC_008519,NC_008582,NC_010107,NC_010226,NC_010329,NC_010739,NC_010817,NC_011051,NC_011109,NC_011280,NC_011530,NC_011765,NC_012123,NC_012213,NC_012485,NC_012486,NC_013035,NC_013117,NC_013237,NC_014143,NC_014185,NC_014326,NC_014469,NC_014952,NC_014953,NC_014954,NC_014955,NC_014956,NC_015267,NC_015268,NC_015325,NC_015691,NC_015692,NC_016013,NC_016014,NC_016074,NC_016075,NC_016157,NC_016898,NC_017716,NC_017862,NC_017993,NC_017994,NC_017995,NC_017996,NC_017997,NC_018074,NC_018075,NC_018076,NC_018575,NC_019023,NC_019852,NC_020084,NC_020085,NC_020500,NC_020501,NC_021472,NC_021483,NC_021930,NC_022095,NC_022253,NC_022373,NC_022647,NC_022892,NC_023178,NC_023496,NC_023852,NC_023873,NC_023882,NC_023891,NC_023894,NC_023895,NC_024300,NC_024893,NC_026640,NC_026946,NC_027528,NC_027779,NC_028125,NC_028126,NC_028267,NC_028492,NC_030151,NC_030795,NC_030796,NC_030797,NC_030798,NC_030799,NC_030800,NC_030801,NC_030839,NC_031756,NC_033740,NC_033745,NC_033781,NC_034616,NC_035193,NC_035199,NC_035201,NC_035208,NC_035478,NC_035479,NC_037059,NC_037061,NC_037064,NC_037067,NC_037069,NC_038516,NC_038517,NC_038518,NC_038519,NC_038520,NC_038521,NC_038522,NC_038523,NC_038524,NC_038525,NC_038526,NC_038527,NC_038531,NC_038889,NC_038914,NC_039036,NC_039037,NC_039038,NC_039039,NC_039040,NC_039041,NC_039042,NC_039086,NC_039089,NC_040548,NC_040550,NC_040569,NC_040578,NC_040579,NC_040580,NC_040583,NC_040604,NC_040619,NC_040620,NC_040640,NC_040655,NC_040688,NC_040691,NC_040709,NC_040727,NC_040728,NC_040785,NC_040787,NC_040803,NC_040804,NC_040805,NC_040806,NC_040818,NC_040827,U06714,U21941,U31778,U31779,U31780,U31781,U31782,U31783,U31784,U31785,U31786,U31787,U31788,U31791,U31793,U31794,U37537,U83595,X05015,X05817,X55964,X55965,X70829,X74462,X74466,X74467,X74468,X74469,X74470,X74471,X74473,X74478,X74479,X74481,X74483,X77858 |

| |
|---|
| **Parvoviridae** |
| AF028704,AF028705,DQ196318,DQ196319,DQ335246,DQ778300,DQ898166,EF441262,EF584447,FJ375127,FJ375128,FJ375129,FJ440683,FJ441297,FJ445512,GQ368252,HM053694,JF429836,JN681175,JQ268283,JQ268284,JX645345,JX827169,KC580640,KF170373,KF214638,KF214640,KF214645,KJ486491,KJ634207,KM105951,KM390024,KM598414,KM598419,KP280068,M81888,NC_000883,NC_000936,NC_001401,NC_001510,NC_001539,NC_001540,NC_001662,NC_001701,NC_001718,NC_001729,NC_001829,NC_001899,NC_002077,NC_002190,NC_003346,NC_004284,NC_004285,NC_004286,NC_004287,NC_004288,NC_004289,NC_004290,NC_004295,NC_004442,NC_004828,NC_005040,NC_005041,NC_005341,NC_005889,NC_006147,NC_006148,NC_006152,NC_006259,NC_006260,NC_006261,NC_006263,NC_006555,NC_007018,NC_007218,NC_007455,NC_011317,NC_011545,NC_012042,NC_012564,NC_012636,NC_012685,NC_012729,NC_014357,NC_014358,NC_014468,NC_014665,NC_015115,NC_015718,NC_016031,NC_016032,NC_016647,NC_016744,NC_016752,NC_017823,NC_018399,NC_018450,NC_019492,NC_020499,NC_022089,NC_022104,NC_022564,NC_022748,NC_022800,NC_023020,NC_023673,NC_023842,NC_023860,NC_024452,NC_024453,NC_024454,NC_024888,NC_025825,NC_025891,NC_025965,NC_026251,NC_026815,NC_026943,NC_027429,NC_028136,NC_028650,NC_028973,NC_029133,NC_029300,NC_029797,NC_030296,NC_030402,NC_030837,NC_030873,NC_031450,NC_031670,NC_031695,NC_031751,NC_031959,NC_032097,NC_034445,NC_034532,NC_035180,NC_035185,NC_035186,NC_037053,NC_038532,NC_038533,NC_038534,NC_038535,NC_038536,NC_038537,NC_038538,NC_038539,NC_038540,NC_038541,NC_038542,NC_038543,NC_038544,NC_038545,NC_038546,NC_038547,NC_038883,NC_038895,NC_038898,NC_039043,NC_039044,NC_039045,NC_039046,NC_039047,NC_039048,NC_039049,NC_039050,NC_040533,NC_040562,NC_040603,NC_040623,NC_040626,NC_040652,NC_040671,NC_040672,NC_040694,NC_040695,NC_040843,NC_043446,U12469,X01457 |
| **Polydnaviridae** |
| AY651828,AY651829,AY651830,DQ075354,DQ075355,DQ075356,DQ075357,DQ075358,DQ075359,DQ075360,EF067319,EF067320,EF067321,EF067322,EF067323,EF067324,EF067325,EF067326,EF067327,EF067328,EF067329,EF067330,EF067331,EF067332,NC_005165,NC_006633,NC_006634,NC_006635,NC_006636,NC_006637,NC_006638,NC_006639,NC_006640,NC_006641,NC_006642,NC_006643,NC_006644,NC_006645,NC_006646,NC_006647,NC_006648,NC_006649,NC_006650,NC_006651,NC_006652,NC_006653,NC_006654,NC_006655,NC_006656,NC_006657,NC_006658,NC_006659,NC_006660,NC_006661,NC_006662,NC_007028,NC_007029,NC_007030,NC_007031,NC_007032,NC_007033,NC_007034,NC_007035,NC_007036,NC_007037,NC_007038,NC_007039,NC_007040,NC_007041,NC_007044,NC_007985,NC_007986,NC_007987,NC_007988,NC_007989,NC_007990,NC_007991,NC_007992,NC_007993,NC_007994,NC_007995,NC_007996,NC_007998,NC_007999,NC_008000,NC_008001,NC_008002,NC_008003,NC_008004,NC_008005,NC_008006,NC_008007,NC_008008,NC_008847,NC_008848,NC_008849,NC_008850,NC_008851,NC_008852,NC_008853,NC_008854,NC_008855,NC_008856,NC_008857,NC_008858,NC_008859,NC_008860,NC_008861,NC_008862,NC_008863,NC_008864,NC_008865,NC_008866,NC_008867,NC_008868,NC_008869,NC_008870,NC_008871,NC_008872,NC_008873,NC_008874,NC_008875,NC_008876,NC_008877,NC_008878,NC_008879,NC_008880,NC_008881,NC_008882,NC_008883,NC_008884,NC_008885,NC_008886,NC_008887,NC_008888,NC_008889,NC_008890,NC_008891,NC_008892,NC_008893,NC_008894,NC_008895,NC_008896,NC_008897,NC_008898,NC_008899,NC_008900,NC_008901,NC_008902,NC_008903,NC_008904,NC_008905,NC_008906,NC_008907,NC_008908,NC_008909,NC_008910,NC_008911,NC_008912,NC_008913,NC_008914,NC_008915,NC_008916,NC_008917,NC_008918,NC_008919,NC_008920,NC_008921,NC_008922,NC_008923,NC_008924,NC_008925,NC_008926,NC_008927,NC_008928,NC_008929,NC_008930,NC_008931,NC_008932,NC_008933,NC_008934,NC_008935,NC_008936,NC_008937,NC_008938,NC_008939,NC_008940,NC_008941,NC_008946,NC_008947,NC_008948,NC_008949,NC_008950,NC_008951,NC_008952,NC_008953,NC_008954,NC_008955,NC_008956,NC_008957,NC_008958,NC_008959,NC_008960,NC_008961,NC_008962,NC_008963,NC_008964,NC_008965,NC_008966,NC_008967,NC_008968,NC_008969,NC_008970,NC_008971,NC_008972,NC_008973,NC_008976,NC_008977,NC_008978,NC_008979,NC_008980,NC_008981,NC_008982,NC_008983,NC_008984,NC_008985,NC_008986,NC_008987,NC_008988,NC_008989,NC_008990,NC_008991,NC_008992,NC_008993,NC_008994,NC_008995,NC_008996,NC_008997,NC_008998,NC_008999,NC_009000,NC_009001,NC_009002,NC_009003,NC_043261,NC_043262,NC_043263,NC_043264,NC_043266,NC_043267,NC_043270,NC_043271,NC_043273,NC_043307,NC_043308,NC_043309,NC_043310,NC_043311,NC_043312,NC_043315,NC_043316,NC_043318,NC_043319,NC_043320,NC_043321,NC_043322,NC_043323,NC_043324,NC_043325,NC_043326,NC_043327,NC_043328,NC_043329,NC_043330,NC_043331,NC_043332,NC_043333,NC_043334,NC_043335,NC_043336,NC_043337,NC_043 |

| |
|---|
| 338,NC_043339,NC_043340,NC_043341,NC_043342,NC_043343,NC_043344,NC_043345,NC_043346,NC_043347,NC_043348,NC_043349,NC_043350,NC_043351,NC_043352,NC_043354,NC_043356,NC_043357,NC_043358,NC_043359,NC_043360,NC_043361,NC_043362 |
| **Polyomaviridae** |
| AB767295,AF118150,DQ192570,DQ192571,EF127906,EF127907,EF127908,FR823284,HG764413,HQ385747,HQ385750,J02288,JX259273,JX262162,KJ577598,KM496323,KM496324,KM496325,M30540,NC_001442,NC_001505,NC_001515,NC_001538,NC_001663,NC_001669,NC_001699,NC_004763,NC_004764,NC_004800,NC_007611,NC_007922,NC_007923,NC_009238,NC_009539,NC_009951,NC_010277,NC_011310,NC_013439,NC_013796,NC_014361,NC_014406,NC_014407,NC_014743,NC_015150,NC_017085,NC_017982,NC_018102,NC_019844,NC_019850,NC_019851,NC_019853,NC_019854,NC_019855,NC_019856,NC_019857,NC_019858,NC_020065,NC_020066,NC_020067,NC_020068,NC_020069,NC_020070,NC_020071,NC_020106,NC_020890,NC_022519,NC_023008,NC_023845,NC_024118,NC_025259,NC_025368,NC_025370,NC_025380,NC_025790,NC_025800,NC_025811,NC_025892,NC_025894,NC_025895,NC_025896,NC_025898,NC_025899,NC_026012,NC_026015,NC_026141,NC_026244,NC_026473,NC_026762,NC_026766,NC_026767,NC_026768,NC_026769,NC_026770,NC_026942,NC_026944,NC_027531,NC_027532,NC_028117,NC_028119,NC_028120,NC_028121,NC_028122,NC_028123,NC_028127,NC_028635,NC_030148,NC_030838,NC_031757,NC_032005,NC_032120,NC_033737,NC_034218,NC_034219,NC_034220,NC_034221,NC_034251,NC_034253,NC_034378,NC_034456,NC_035181,NC_038554,NC_038555,NC_038556,NC_038557,NC_038558,NC_038559,NC_039051,NC_039052,NC_039053,NC_040538,NC_040566,NC_040573,NC_040598,NC_040600,NC_040607,NC_040634,NC_040638,NC_040676,NC_040677,NC_040705,NC_040714,NC_040715,NC_040821,NC_040822 |
| **Riboviria** |
| AB032553,AB042808,AB050936,AB073912,AB090161,AB187514,AB194796,AB205396,AB220921,AB252582,AB365435,AB426611,AB447427,AB447428,AB447429,AB447430,AB447431,AB447432,AB447433,AB447434,AB447435,AB447436,AB447437,AB447438,AB447439,AB447440,AB447441,AB447442,AB447443,AB447444,AB447445,AB447446,AB447447,AB447448,AB447449,AB447450,AB447451,AB447452,AB447453,AB447454,AB447455,AB447456,AB447457,AB447458,AB447459,AB447460,AB447461,AB447462,AB447463,AB541201,AB541202,AB541203,AB541204,AB541205,AB543808,AB558119,AB593690,AB614356,AB678778,AB690461,AB795432,AC_000192,AF002227,AF039205,AF046869,AF057136,AF059242,AF059243,AF070476,AF079457,AF081485,AF083069,AF086833,AF091605,AF091736,AF093797,AF103734,AF123432,AF123433,AF145896,AF162711,AF201929,AF227250,AF230973,AF241359,AF260508,AF274010,AF309418,AF311056,AF311938,AF311939,AF316321,AF326963,AF327920,AF327921,AF327922,AF338106,AF352027,AF361253,AF389115,AF389116,AF389117,AF389452,AF389453,AF389454,AF389455,AF389456,AF389462,AF389463,AF389464,AF389465,AF389466,AF407339,AF457102,AF524867,AF525933,AJ005695,AJ132961,AJ132997,AJ276479,AJ276480,AJ276481,AJ577589,AJ781401,AJ880277,AJ889866,AJ889867,AJ889868,AJ889918,AM113988,AM157175,AM235750,AM404308,AM498051,AM498052,AM498053,AM744987,AM744988,AM744989,AM744997,AM744998,AM744999,AM745007,AM745008,AM745009,AM745017,AM745018,AM745019,AM745027,AM745028,AM745029,AM745035,AM745037,AM745038,AM745039,AM745047,AM745048,AM745049,AM745057,AM745058,AM745059,AM745067,AM745068,AM745069,AM745077,AM745078,AM745079,AM910652,AY010722,AY032605,AY134748,AY260942,AY260943,AY260944,AY260949,AY260950,AY260951,AY278488,AY278491,AY278554,AY278741,AY297819,AY302539,AY302540,AY302541,AY302542,AY302543,AY302544,AY302545,AY302546,AY302547,AY302548,AY302549,AY302550,AY302551,AY302552,AY302553,AY302554,AY302555,AY302556,AY302557,AY302559,AY302560,AY350750,AY353550,AY357075,AY357076,AY394850,AY429470,AY462107,AY485642,AY486084,AY508697,AY515512,AY518894,AY554397,AY556057,AY556070,AY575773,AY588319,AY593765,AY593796,AY593805,AY593806,AY593808,AY593809,AY593840,AY593847,AY593851,AY646283,AY646511,AY685920,AY685921,AY686687,AY729016,AY741811,AY743910,AY751783,AY772730,AY773285,AY800279,AY842931,AY843297,AY843298,AY843299,AY843300,AY843301,AY843302,AY843303,AY843304,AY843305,AY843306,AY843307,AY843308,AY859526,AY863002,AY864805,AY864806,AY876912,AY876913,AY898809,CY011117,CY011118,CY011119,CY011125,CY011126,CY011127,CY011133,CY011134,CY011135,CY011141,CY011142,CY011143,D00239,D00435,D00507,D00538,D00627,D00820,D13096,D90457,DQ011234,DQ011855,DQ028633,DQ058829,DQ070852,DQ217792,DQ238861,DQ256132,DQ256133,DQ256134,DQ286292,DQ294633,DQ315670,DQ328874,DQ328875,DQ358078,DQ369797,DQ399290,DQ412042,DQ412043,DQ447649,DQ447652,DQ447657,DQ456824,DQ473486,DQ473488,DQ473489,DQ473490,DQ473491,DQ473492,DQ473493,DQ473494,DQ473497,DQ473499,DQ473500,DQ473504,DQ473505,DQ473506,DQ473507,DQ473508,DQ473510,DQ473511,DQ480514,DQ640652,DQ648794, |

| |
|---|
| DQ648856,DQ648857,DQ658413,DQ811787,DQ812092,DQ812093,DQ848678,DQ851494,DQ902712,DQ902713,DQ911368,DQ915164,DQ995634,DQ995640,DQ995647,EF011023,EF014462,EF015886,EF017707,EF065505,EF065506,EF065507,EF065508,EF065509,EF065510,EF065511,EF065512,EF065513,EF065514,EF065515,EF065516,EF067923,EF067924,EF107097,EF108464,EF173414,EF173415,EF173420,EF173423,EF173425,EF424615,EF424616,EF424617,EF424618,EF424619,EF424620,EF424621,EF424622,EF424623,EF424624,EF424625,EF424626,EF424627,EF424628,EF424629,EF429197,EF429198,EF429199,EF429200,EF446132,EF446615,EF552688,EF552689,EF552690,EF552691,EF552692,EF552693,EF552694,EF552695,EF552696,EF552697,EF555644,EF555645,EF558545,EF667343,EF667344,EU004663,EU004664,EU004665,EU004666,EU004667,EU004668,EU004669,EU004670,EU004671,EU004672,EU004673,EU004674,EU004675,EU004676,EU004677,EU004678,EU004679,EU004680,EU004681,EU004682,EU004683,EU020009,EU037962,EU140838,EU143843,EU155216,EU155260,EU371559,EU371560,EU371561,EU371562,EU371563,EU371564,EU420137,EU420138,EU439428,EU563512,EU627591,EU716175,EU755009,EU779803,EU815052,EU854589,FJ009367,FJ355929,FJ355930,FJ376620,FJ387164,FJ415324,FJ425184,FJ425185,FJ425186,FJ425187,FJ425188,FJ425189,FJ434664,FJ445112,FJ445113,FJ445114,FJ445116,FJ445118,FJ445119,FJ445120,FJ445121,FJ445122,FJ445123,FJ445124,FJ445125,FJ445126,FJ445127,FJ445128,FJ445129,FJ445130,FJ445131,FJ445132,FJ445133,FJ445134,FJ445135,FJ445136,FJ445138,FJ445140,FJ445141,FJ445142,FJ445143,FJ445144,FJ445145,FJ445146,FJ445147,FJ445148,FJ445149,FJ445150,FJ445151,FJ445152,FJ445153,FJ445154,FJ445155,FJ445156,FJ445157 |
| **Test-2; Source: Virus-Host-DB** |
| **Betaflexiviridae** |
| AF057136,NC_001946,NC_038324,NC_038325,NC_038966,NC_039087,NC_040545,NC_040554,NC_040564,NC_040568,NC_040616,NC_040627,NC_001948,NC_040630,NC_040643,NC_040689,NC_040703,NC_040797,NC_040800,NC_043081,NC_043082,NC_043086,NC_043087,NC_002468,NC_043088,NC_043412,NC_002500,NC_002552,NC_002729,NC_002795,NC_003462,NC_003499,NC_003557,AY646511,NC_003602,NC_003604,NC_003689,NC_003870,NC_003877,NC_005138,NC_005343,NC_006550,NC_006946,NC_007289,EU020009,NC_008020,NC_008266,NC_008292,NC_008552,NC_009087,NC_009383,NC_009759,NC_009764,NC_009892,NC_009991,FJ009367,NC_010305,NC_010538,NC_011062,NC_011106,NC_011525,NC_011540,NC_011552,NC_012038,NC_012210,NC_012519,JF320811,NC_012869,NC_013006,NC_013527,NC_014730,NC_014821,NC_015220,NC_015395,NC_015782,NC_016080,NC_016404,JX559646,NC_016440,NC_017859,NC_018175,NC_018448,NC_018458,NC_018714,NC_019025,NC_019029,NC_019030,NC_020996,NC_001361,NC_023295,NC_023892,NC_024449,NC_024686,NC_025388,NC_025468,NC_025469,NC_026248,NC_026616,NC_027527,NC_001409,NC_028111,NC_028868,NC_028975,NC_029085,NC_029086,NC_029087,NC_029088,NC_029089,NC_029301,NC_030657,NC_001749,NC_030926,NC_031089,NC_034264,NC_034376,NC_034377,NC_034833,NC_035202,NC_035203,NC_035462,NC_037058 |
| **Bromoviridae** |
| AJ276479,AJ276480,AJ276481,NC_001440,NC_001495,NC_002024,NC_002025,NC_002026,NC_002027,NC_002028,NC_002034,NC_002035,NC_002038,NC_002039,NC_002040,NC_003451,NC_003452,NC_003453,NC_003464,NC_003465,NC_003480,NC_003541,NC_003542,NC_003543,NC_003546,NC_003547,NC_003548,NC_003568,NC_003569,NC_003570,NC_003649,NC_003650,NC_003651,NC_003671,NC_003673,NC_003674,NC_003808,NC_003809,NC_003810,NC_003833,NC_003834,NC_003835,NC_003836,NC_003837,NC_003838,NC_003842,NC_003844,NC_003845,NC_004006,NC_004007,NC_004008,NC_004120,NC_004121,NC_004122,NC_004362,NC_004363,NC_005848,NC_005849,NC_005854,NC_006064,NC_006065,NC_006566,NC_006567,NC_006568,NC_006999,NC_007000,NC_007001,NC_008037,NC_008038,NC_008039,NC_008706,NC_008707,NC_008708,NC_009536,NC_009537,NC_009538,NC_011553,NC_011554,NC_011555,NC_011807,NC_011808,NC_011809,NC_012134,NC_012135,NC_012136,NC_013266,NC_013267,NC_013268,NC_018402,NC_018403,NC_018404,NC_022127,NC_022128,NC_022129,NC_022250,NC_022251,NC_022252,NC_025477,NC_025478,NC_025481,NC_025482,NC_025483,NC_025484,NC_027928,NC_027929,NC_027930,NC_038776,NC_038777,NC_039074,NC_039075,NC_039076,NC_040389,NC_040390,NC_040391,NC_040392,NC_040393,NC_040394,NC_040435,NC_040436,NC_040437,NC_040469,NC_040471 |
| **Caliciviridae** |
| AB042808,AB187514,AB220921,AB365435,AB447427,AB447428,AB447429,AB447430,AB447431,AB447432,AB447433,AB447434,AB447435,AB447436,AB447437,AB447438,AB447439,AB447440,AB447441,AB447442,AB447443,AB447444,AB447445,AB447446,AB447447,AB447448,AB447449,AB447450 |

| |
|---|
| ,AB447451,AB447452,AB447453,AB447454,AB447455,AB447456,AB447457,AB447458,AB447459,AB447460,AB447461,AB447462,AB447463,AB541201,AB541202,AB541203,AB541204,AB541205,AB543808,AB614356,AF091736,AF093797,AF145896,AY032605,AY134748,AY485642,AY741811,AY772730,DQ058829,DQ369797,DQ456824,DQ658413,DQ911368,EF014462,EU004663,EU004664,EU004665,EU004666,EU004667,EU004668,EU004669,EU004670,EU004671,EU004672,EU004673,EU004674,EU004675,EU004676,EU004677,EU004678,EU004679,EU004680,EU004681,EU004682,EU004683,EU854589,FJ355929,FJ355930,FJ387164,FJ514242,FJ515294,FJ537135,FJ537136,FJ537137,FJ537138,GQ475301,GQ475302,GU594162,GU980585,GU991353,GU991354,GU991355,HF952119,HF952120,HF952121,HF952122,HF952123,HF952124,HF952125,HF952126,HF952127,HF952128,HF952129,HF952130,HF952131,HF952132,HF952133,HF952134,HF952135,HM002617,HQ009513,HQ392821,HQ449728,HQ664990,JF320644,JF320645,JF320646,JF320647,JF320648,JF320649,JF320650,JF320651,JF320652,JF320653,JF781268,JN400599,JN400600,JN400601,JN400602,JN400603,JN400604,JN400605,JN400606,JN400607,JN400608,JN400609,JN400610,JN400611,JN400612,JN400613,JN400614,JN400615,JN400616,JN400617,JN400618,JN400619,JN400620,JN400621,JN400622,JN400623,JN400624,JN400625,JN400626,JN595867,JQ388274,JQ613567,JQ613568,JQ613569,JQ613570,JQ622197,JQ798158,JQ911594,JQ911595,JQ911596,JQ911597,JQ911598,JX018212,JX023285,JX023286,JX047864,JX126912,JX126913,JX439815,JX439816,JX439817,JX439818,JX439819,JX448566,JX459900,JX459901,JX459902,JX459903,JX459904,JX459905,JX459906,JX459907,JX459908,JX846924,JX846927,JX989073,JX989074,JX989075,JX993277,KC013592,KC175323,KC175342,KC175343,KC175344,KC175345,KC175346,KC175347,KC175348,KC175349,KC175350,KC175351,KC175352,KC175353,KC175354,KC175355,KC175356,KC175357,KC175358,KC175359,KC175360,KC175361,KC175362,KC175363,KC175364,KC175365,KC175366,KC175367,KC175368,KC175369,KC175370,KC175371,KC175372,KC175373,KC175374,KC175375,KC175376,KC175377,KC175378,KC175379,KC175380,KC175381,KC175382,KC175383,KC175384,KC175385,KC175386,KC175387,KC175388,KC175389,KC175390,KC175391,KC175392,KC175393,KC175394,KC175395,KC175396,KC175397,KC175398,KC175399,KC175400,KC175401,KC175402,KC175403,KC175404,KC175405,KC175406,KC175407,KC175408,KC175409,KC175410,KC409301,KC409302,KC463910,KC464496,KC464497,KC464498,KC464499,KC464500,KC577174,KC577175,KC631827,KC792553,KC894731,KC894942,KC894943,KC960615,KF204570,KF306212,KF306213,KF306214,KF429760,KF429761,KF429765,KF429766,KF429768,KF429770,KF429773,KF429774,KF429776,KF429777,KF429778,KF429783,KF429787,KF429789,KF429790,KF712491,KF712496,KF712497,KF712498,KF712499,KF712501,KF712502,KF712504,KF712510,KJ196276,KJ196277,KJ196278,KJ196279,KJ196280,KJ196281,KJ196282,KJ196283,KJ196284,KJ196285,KJ196286,KJ196287,KJ196288,KJ196289,KJ196293,KJ196294,KJ196295,KJ196296,KJ196297,KJ196298,KJ196299,KJ407072,KJ407073,KJ407074,KJ407075,KJ407076,KJ508818,KJ541743,KJ649705,KJ685403,KJ685405,KJ685408,KJ685412,KJ685413,KJ685414,KJ685415,KJ685417,KM272334,NC_000940,NC_001481,NC_001543,NC_001959,NC_002551,NC_002615,NC_004064,NC_004541,NC_004542,NC_006269,NC_006554,NC_006875,NC_007916,NC_008311,NC_008580,NC_010624,NC_011050,NC_011704,NC_012699,NC_017936,NC_019712,NC_024031,NC_024078,NC_025676,NC_027026,NC_027122,NC_029645,NC_029646,NC_029647,NC_030793,NC_031324,NC_033081,NC_033776,NC_034444,NC_035675,NC_039475,NC_039476,NC_039477,NC_039897,NC_040674,NC_040876,NC_043512,NC_043516,NC_044045,NC_044046,NC_044047,U15301,U54983,X86557 |
| **Coronaviridae** |
| MG772933.1,MG772934.1,AC_000192,AF201929,AY278488,AY278491,AY278554,AY278741,AY350750,AY357075,AY357076,AY394850,AY515512,AY518894,AY646283,AY864805,AY864806,D13096,DQ011855,DQ412042,DQ412043,DQ640652,DQ648794,DQ648856,DQ648857,DQ811787,DQ848678,DQ915164,EF065505,EF065506,EF065507,EF065508,EF065509,EF065510,EF065511,EF065512,EF065513,EF065514,EF065515,EF065516,EF424615,EF424616,EF424617,EF424618,EF424619,EF424620,EF424621,EF424622,EF424623,EF424624,EF446615,EU371559,EU371560,EU371561,EU371562,EU371563,EU371564,EU420137,EU420138,FJ376620,FJ415324,FJ425184,FJ425185,FJ425186,FJ425187,FJ425188,FJ425189,FJ647218,FJ647219,FJ647220,FJ647221,FJ647222,FJ647223,FJ647224,FJ647225,FJ647226,FJ647227,FJ882935,FJ882942,FJ882945,FJ882954,FJ882963,FJ884686,FJ938051,FJ938052,FJ938053,FJ938054,FJ938055,FJ938056,FJ938057,FJ938058,FJ938059,FJ938060,FJ938061,FJ938062,FJ938063,FJ938064,FJ938065,FJ938066,FJ938067,FN430414,FN430415,GQ153539,GQ153540,GQ153541,GQ153542,GQ153543,GQ153544,GQ153545,GQ153546,GQ153547,GQ153548,GU553361,GU553362,HM211098,HM211099,HM211100,HM211101,HM245926,HQ392469,HQ392470,HQ392471,HQ392472,JF705860,JF792616,JN183882,JN183883,JQ173883,JQ410000,JQ989272,JX169867,JX860640,JX869059,JX993987,JX993988,KC667074,KC776174,KC881005,KC881006,KF367457,KF569996,KF793824,KF906249,KJ473821,KJ481931,KJ567050,KJ601777,KJ601778,KJ601779,KJ601780, |

<table>
<tr><td>KJ769231,KM820765,KP981395,LM645057,LN610099,NC_001451,NC_001846,NC_002306,NC_002645,NC_003045,NC_003436,NC_004718,NC_005831,NC_006213,NC_006577,NC_009019,NC_009020,NC_009021,NC_009657,NC_009988,NC_010437,NC_010438,NC_010646,NC_010800,NC_011547,NC_011549,NC_011550,NC_012936,NC_014470,NC_016991,NC_016992,NC_016993,NC_016994,NC_016995,NC_016996,NC_017083,NC_018871,NC_019843,NC_022103,NC_023760,NC_025217,NC_026011,NC_028752,NC_028806,NC_028811,NC_028814,NC_028824,NC_028833,NC_030292,NC_030886,NC_032107,NC_032730,NC_034440,NC_034972,NC_035191,NC_038294,NC_038861,NC_039207,NC_039208,BetaCoV/bat/Yunnan/RaTG13/2013|EPI_ISL_402131</td></tr>
<tr><td>Flaviviridae</td></tr>
<tr><td>NC_027819,NC_027998,NC_027999,NC_028137,NC_028377,NC_029054,NC_029055,NC_030289,NC_030290,NC_030291,NC_030400,NC_030401,NC_030653,NC_030791,NC_031327,NC_031916,NC_031947,NC_031950,NC_032088,NC_033693,NC_033694,NC_033697,NC_033698,NC_033699,NC_033715,NC_033721,NC_033723,NC_033724,NC_033725,NC_033726,NC_034007,NC_034017,NC_034018,NC_034151,NC_034204,NC_034222,NC_034223,NC_034224,NC_034225,NC_034242,NC_034442,NC_035071,NC_035118,NC_035187,NC_035432,NC_035889,NC_038425,NC_038426,NC_038427,NC_038428,NC_038429,NC_038430,NC_038431,NC_038432,NC_038433,NC_038434,NC_038435,NC_038436,NC_038437,NC_038882,NC_038912,NC_038964,NC_039218,NC_039219,NC_039237,NC_040555,NC_040589,NC_040610,NC_040645,NC_040682,NC_040776,NC_040788,NC_040815,NC_043110,U70263,Z46258,AB690461,AB795432,AF002227,AF070476,AF091605,AF311056,AF326963,AF407339,AJ132997,AM404308,AM910652,AY554397,AY842931,AY859526,AY863002,AY898809,DQ480514,EF424625,EF424626,EF424627,EF424628,EF424629,EF429197,EF429198,EF429199,EF429200,EU155216,EU155260,FJ654700,GQ275355,HQ231763,JN704144,JN860200,JQ289550,JQ920421,JX196334,JX227952,JX227953,JX227954,JX227955,JX227958,JX227960,JX227962,JX227963,JX227965,JX227967,JX227970,JX227972,JX227979,JX477686,KC815310,KC815311,KC990542,KF907503,KF917538,KJ469370,KJ660072,KM225263,KM225264,KM225265,KM408491,M91671,NC_000943,NC_001437,NC_001461,NC_001474,NC_001475,NC_001477,NC_001563,NC_001564,NC_001655,NC_001672,NC_001710,NC_001809,NC_001837,NC_002031,NC_002640,NC_002657,NC_003635,NC_003675,NC_003676,NC_003678,NC_003679,NC_003687,NC_003690,NC_003996,NC_004102,NC_004119,NC_005039,NC_005062,NC_005064,NC_006551,NC_007580,NC_008604,NC_008718,NC_008719,NC_009026,NC_009028,NC_009029,NC_009823,NC_009824,NC_009825,NC_009826,NC_009827,NC_009942,NC_012532,NC_012533,NC_012534,NC_012671,NC_012735,NC_012812,NC_012932,NC_015843,NC_016997,NC_017086,NC_018705,NC_018713,NC_020902,NC_021069,NC_021153,NC_021154,NC_023176,NC_023424,NC_023439,NC_024017,NC_024018,NC_024077,NC_024111,NC_024112,NC_024113,NC_024114,NC_024299,NC_024377,NC_024805,NC_024806,NC_024889,NC_025672,NC_025673,NC_025677,NC_025679,NC_026620,NC_026623,NC_026624,NC_026797,NC_027709,NC_027817</td></tr>
<tr><td>Peribunyaviridae</td></tr>
<tr><td>NC_001925,NC_001926,NC_004108,NC_004109,NC_005775,NC_005776,NC_009894,NC_009895,NC_018459,NC_018461,NC_018463,NC_018465,NC_018466,NC_018467,NC_018476,NC_018478,NC_021242,NC_021243,NC_022038,NC_022039,NC_022595,NC_022596,NC_024074,NC_024076,NC_026281,NC_026283,NC_026617,NC_026618,NC_026619,NC_027715,NC_027717,NC_031135,NC_031136,NC_031221,NC_031222,NC_031287,NC_031288,NC_031291,NC_031292,NC_034459,NC_034460,NC_034461,NC_034468,NC_034475,NC_034477,NC_034479,NC_034482,NC_034487,NC_034488,NC_034489,NC_034490,NC_034491,NC_034492,NC_034493,NC_034495,NC_034497,NC_034499,NC_034500,NC_034504,NC_034505,NC_034506,NC_034631,NC_034633,NC_038713,NC_038714,NC_038715,NC_038717,NC_038718,NC_038720,NC_038723,NC_038724,NC_038727,NC_038728,NC_038729,NC_038730,NC_038733,NC_038734,NC_038735,NC_038736,NC_038738,NC_038739,NC_038741,NC_038742,NC_038942,NC_039183,NC_039184,NC_039186,NC_039187,NC_043036,NC_043037,NC_043546,NC_043548,NC_043550,NC_043551,NC_043552,NC_043553,NC_043555,NC_043556,NC_043559,NC_043560,NC_043561,NC_043563,NC_043564,NC_043565,NC_043567,NC_043568,NC_043570,NC_043571,NC_043573,NC_043575,NC_043577,NC_043578,NC_043579,NC_043580,NC_043583,NC_043584,NC_043586,NC_043587,NC_043588,NC_043589,NC_043591,NC_043592,NC_043594,NC_043595,NC_043597,NC_043599,NC_043600,NC_043602,NC_043603,NC_043605,NC_043607,NC_043608,NC_043612,NC_043614,NC_043615,NC_043617,NC_043618,NC_043619,NC_043621,NC_043623,NC_043627,NC_043629,NC_043630,NC_043632,NC_043633,NC_043634,NC_043637,NC_043638,NC_043639,NC_043641,NC_043645,NC_043646,NC_043651,NC_043652,NC_043653,NC_043655,NC_043674,NC_043675,NC_043687,NC_043688,NC_043690,NC_043691,NC_043692,NC_043694,NC_043697,NC_043699</td></tr>
</table>

| Phenuiviridae |
| --- |
| NC_002323,NC_002324,NC_002325,NC_002326,NC_002327,NC_002328,NC_003753,NC_003754,NC_003755,NC_003776,NC_005214,NC_005220,NC_006319,NC_006320,NC_014396,NC_014397,NC_015373,NC_015374,NC_015411,NC_015412,NC_015450,NC_015451,NC_018136,NC_018138,NC_022630,NC_022631,NC_023633,NC_023635,NC_024494,NC_024495,NC_027140,NC_027141,NC_029082,NC_029127,NC_029128,NC_029901,NC_029903,NC_031138,NC_031139,NC_031295,NC_031298,NC_031313,NC_031316,NC_031317,NC_031318,NC_031320,NC_031321,NC_032158,NC_032159,NC_032257,NC_032276,NC_032277,NC_032278,NC_032280,NC_032282,NC_033830,NC_033835,NC_033836,NC_033838,NC_033840,NC_033841,NC_033842,NC_033844,NC_033846,NC_033847,NC_033848,NC_033849,NC_036597,NC_036598,NC_036602,NC_036604,NC_036605,NC_037612,NC_037614,NC_037616,NC_038257,NC_038258,NC_038261,NC_038262,NC_038748,NC_038750,NC_038751,NC_038752,NC_038754,NC_038757,NC_038934,NC_039191,NC_039192,NC_040450,NC_040493,NC_040494,NC_043045,NC_043046,NC_043049,NC_043051,NC_043450,NC_043451,NC_043477,NC_043481,NC_043482,NC_043509,NC_043510,NC_043609,NC_043611,NC_043679,NC_043680,X89628 |
| **Picornaviridae** |
| AB090161,AB205396,AB252582,AB426611,AB678778,AF039205,AF081485,AF083069,AF123432,AF123433,AF162711,AF230973,AF241359,AF274010,AF311938,AF311939,AF316321,AF327920,AF327921,AF327922,AF352027,AF361253,AF524867,AJ005695,AJ132961,AJ577589,AJ889918,AM235750,AY302539,AY302540,AY302541,AY302542,AY302543,AY302544,AY302545,AY302546,AY302547,AY302548,AY302549,AY302550,AY302551,AY302552,AY302553,AY302554,AY302555,AY302556,AY302557,AY302559,AY302560,AY429470,AY462107,AY508697,AY556057,AY556070,AY593765,AY593796,AY593805,AY593806,AY593808,AY593809,AY593840,AY593847,AY593851,AY686687,AY751783,AY773285,AY843297,AY843298,AY843299,AY843300,AY843301,AY843302,AY843303,AY843304,AY843305,AY843306,AY843307,AY843308,AY876912,AY876913,D00239,D00435,D00538,D00627,D00820,D90457,DQ256132,DQ256133,DQ256134,DQ294633,DQ315670,DQ358078,DQ473486,DQ473488,DQ473489,DQ473490,DQ473491,DQ473492,DQ473493,DQ473494,DQ473497,DQ473499,DQ473500,DQ473504,DQ473505,DQ473506,DQ473507,DQ473508,DQ473510,DQ473511,DQ812092,DQ812093,DQ902712,DQ902713,DQ995634,DQ995640,DQ995647,EF015886,EF067923,EF067924,EF107097,EF173414,EF173415,EF173420,EF173423,EF173425,EF552688,EF552689,EF552690,EF552691,EF552692,EF552693,EF552694,EF552695,EF552696,EF552697,EF555644,EF555645,EF667343,EF667344,EU140838,EU716175,EU755009,EU815052,FJ445112,FJ445113,FJ445114,FJ445116,FJ445118,FJ445119,FJ445120,FJ445121,FJ445122,FJ445123,FJ445124,FJ445125,FJ445126,FJ445127,FJ445128,FJ445129,FJ445130,FJ445131,FJ445132,FJ445133,FJ445134,FJ445135,FJ445136,FJ445138,FJ445140,FJ445141,FJ445142,FJ445143,FJ445144,FJ445145,FJ445146,FJ445147,FJ445148,FJ445149,FJ445150,FJ445151,FJ445152,FJ445153,FJ445154,FJ445155,FJ445156,FJ445157,FJ445160,FJ445161,FJ445162,FJ445163,FJ445164,FJ445165,FJ445167,FJ445168,FJ445169,FJ445170,FJ445171,FJ445172,FJ445173,FJ445174,FJ445175,FJ445176,FJ445178,FJ445179,FJ445180,FJ445181,FJ445182,FJ445183,FJ445185,FJ445186,FJ445187,FJ445188,FJ445189,FJ445190,FM955278,GQ122332,GQ249161,GQ323774,GQ485310,GQ485311,GQ865517,HM185056,HM777023,HQ400942,HQ654774,HQ702854,HQ728260,HQ728261,HQ728262,HQ875059,JF905564,JN088541,JN379039,JN710381,JQ277724,JQ814852,JQ818253,JQ898342,JQ911763,JQ975417,JX050181,JX174177,JX262382,JX491648,JX961709,JX982257,KC663628,KC811837,KF312882,KF422142,KF831027,KF874626,KF958308,KF990476,KJ857508,KM203656,KM609480,KP036483,L24917,LK021688,M12197,M16560,M20301,NC_001366,NC_001430,NC_001472,NC_001479,NC_001489,NC_001490,NC_001612,NC_001617,NC_001859,NC_001897,NC_001918,NC_002058,NC_003976,NC_003983,NC_003985,NC_003987,NC_003988,NC_003990,NC_004421,NC_004441,NC_004451,NC_006553,NC_008250,NC_008714,NC_009448,NC_009891,NC_009996,NC_010354,NC_010415,NC_010810,NC_011349,NC_011829,NC_012798,NC_012800,NC_012801,NC_012802,NC_012957,NC_012986,NC_013695,NC_014411,NC_014412,NC_014413,NC_015626,NC_015934,NC_015936,NC_015940,NC_015941,NC_016156,NC_016403,NC_016769,NC_018226,NC_018400,NC_018506,NC_018668,NC_021178,NC_021201,NC_021220,NC_021482,NC_022332,NC_022802,NC_023162,NC_023422,NC_023858,NC_023861,NC_023984,NC_023985,NC_023987,NC_023988,NC_024070,NC_024073,NC_024120,NC_024765,NC_024766,NC_024767,NC_024768,NC_024769,NC_024770,NC_025114,NC_025432,NC_025474,NC_025675,NC_025890,NC_025961,NC_026249,NC_026314,NC_026315,NC_026316,NC_026470,NC_026921,NC_027054,NC_027214,NC_027818,NC_027918,NC_027919,NC_028363,NC_028364,NC_028365,NC_028366,NC_028380,NC_028964,NC_028970,NC_028981,NC_029854,NC_029905,NC_030454,NC_030843,NC_031105,NC_031106,NC_032126,NC_033695,NC_033793,NC_033818,NC_033819,NC_033820,NC_034206,NC_034245,NC_034267,NC_034381,NC_034385,NC_034453,NC_034617,NC_034971,NC_035110,NC_035198,NC_035779,NC_036 |

| |
|---|
| 588,NC_037654,NC_038303,NC_038304,NC_038305,NC_038306,NC_038307,NC_038308,NC_038309,NC_038310,NC_038311,NC_038312,NC_038313,NC_038314,NC_038315,NC_038316,NC_038317,NC_038318,NC_038319,NC_038878,NC_038880,NC_038957,NC_038961,NC_038989,NC_039004,NC_039209,NC_039210,NC_039211,NC_039212,NC_039235,NC_040605,NC_040611,NC_040642,NC_040673,NC_040684,NC_043071,NC_043072,NC_043544,U05876,U16283,U22521,V01149,X00925,X05690,X56019,X67706,X77708,X84981,X92886 |
| **Potyviridae** |
| AB194796,AJ889866,AJ889867,AJ889868,AM113988,AM157175,AY010722,AY575773,D00507,DQ851494,EF017707,EF558545,EU563512,HE608963,HE608964,HF585099,HF585103,HM590055,JQ924285,JQ924286,NC_000947,NC_001445,NC_001517,NC_001555,NC_001616,NC_001671,NC_001768,NC_001785,NC_001814,NC_001841,NC_001886,NC_002349,NC_002350,NC_002509,NC_002600,NC_002634,NC_002990,NC_002991,NC_003224,NC_003377,NC_003397,NC_003398,NC_003399,NC_003482,NC_003483,NC_003492,NC_003501,NC_003536,NC_003537,NC_003605,NC_003606,NC_003742,NC_003797,NC_004010,NC_004011,NC_004013,NC_004016,NC_004017,NC_004035,NC_004039,NC_004047,NC_004426,NC_004573,NC_004752,NC_005028,NC_005029,NC_005136,NC_005288,NC_005304,NC_005778,NC_005903,NC_005904,NC_006262,NC_006941,NC_007147,NC_007180,NC_007216,NC_007433,NC_007728,NC_007913,NC_008028,NC_008393,NC_008558,NC_008824,NC_009741,NC_009742,NC_009744,NC_009745,NC_009805,NC_009994,NC_009995,NC_010521,NC_010735,NC_010736,NC_010954,NC_011541,NC_011560,NC_011918,NC_012698,NC_012799,NC_013261,NC_014037,NC_014038,NC_014064,NC_014252,NC_014325,NC_014327,NC_014536,NC_014742,NC_014790,NC_014791,NC_014898,NC_014905,NC_015393,NC_015394,NC_016044,NC_016159,NC_016441,NC_017824,NC_017967,NC_017970,NC_017977,NC_018093,NC_018176,NC_018455,NC_018572,NC_018833,NC_018872,NC_019031,NC_019409,NC_019412,NC_019415,NC_020072,NC_020105,NC_020896,NC_021065,NC_021197,NC_021786,NC_022745,NC_023014,NC_023175,NC_023628,NC_024471,NC_025250,NC_025254,NC_025821,NC_026615,NC_026759,NC_027210,NC_027706,NC_028144,NC_028145,NC_029051,NC_029076,NC_030118,NC_030236,NC_030293,NC_030391,NC_030794,NC_030840,NC_030847,NC_031339,NC_032912,NC_034208,NC_034273,NC_034835,NC_035134,NC_035458,NC_035459,NC_035461,NC_036802,NC_037051,NC_038560,NC_038561,NC_038562,NC_038920,NC_038984,NC_039002,NC_039088,NC_040507,NC_040508,NC_040650,NC_040802,NC_040836,NC_043133,NC_043141,NC_043149,NC_043165,NC_043168,NC_043171,NC_043172,NC_043424,NC_043532,NC_043536,NC_043537,U05771 |
| **Reoviridae** |
| AF389452,AF389453,AF389454,AF389455,AF389456,AF389462,AF389463,AF389464,AF389465,AF389466,AM498051,AM498052,AM498053,AM744987,AM744988,AM744989,AM744997,AM744998,AM744999,AM745007,AM745008,AM745009,AM745017,AM745018,AM745019,AM745027,AM745028,AM745029,AM745035,AM745037,AM745038,AM745039,AM745047,AM745048,AM745049,AM745057,AM745058,AM745059,AM745067,AM745068,AM745069,AM745077,AM745078,AM745079,FN563984,HG513046,NC_002557,NC_002558,NC_002559,NC_002560,NC_002567,NC_003006,NC_003007,NC_003008,NC_003009,NC_003010,NC_003016,NC_003017,NC_003018,NC_003019,NC_003020,NC_003654,NC_003655,NC_003656,NC_003657,NC_003658,NC_003659,NC_003696,NC_003697,NC_003698,NC_003699,NC_003700,NC_003701,NC_003702,NC_003703,NC_003728,NC_003729,NC_003730,NC_003734,NC_003735,NC_003736,NC_003737,NC_003749,NC_003750,NC_003751,NC_003752,NC_003759,NC_003761,NC_003762,NC_003771,NC_003772,NC_003773,NC_003774,NC_004181,NC_004182,NC_004183,NC_004184,NC_004185,NC_004186,NC_004187,NC_004188,NC_004210,NC_004211,NC_004212,NC_004213,NC_004214,NC_004217,NC_004218,NC_004219,NC_005166,NC_005167,NC_005168,NC_005169,NC_005170,NC_005171,NC_005986,NC_005989,NC_005990,NC_005996,NC_005997,NC_005998,NC_005999,NC_006000,NC_006013,NC_006014,NC_006017,NC_006021,NC_006023,NC_007154,NC_007155,NC_007157,NC_007158,NC_007159,NC_007160,NC_007163,NC_007524,NC_007525,NC_007533,NC_007534,NC_007535,NC_007536,NC_007546,NC_007547,NC_007548,NC_007549,NC_007550,NC_007551,NC_007559,NC_007560,NC_007561,NC_007562,NC_007563,NC_007572,NC_007574,NC_007582,NC_007583,NC_007584,NC_007586,NC_007592,NC_007656,NC_007657,NC_007658,NC_007666,NC_007667,NC_007668,NC_007669,NC_007670,NC_007736,NC_007737,NC_007738,NC_007739,NC_007748,NC_007749,NC_007750,NC_008171,NC_008172,NC_008173,NC_008174,NC_008175,NC_008729,NC_008730,NC_008731,NC_008732,NC_008733,NC_008735,NC_008736,NC_009243,NC_009244,NC_009247,NC_009248,NC_009249,NC_010584,NC_010585,NC_010586,NC_010587,NC_010588,NC_010589,NC_010666,NC_010667,NC_010668,NC_010669,NC_010670,NC_010743,NC_010744,NC_010745,NC_010746,NC_010747,NC_010748,NC_011506,NC_011507,NC_011508,NC_011510,NC_012535,NC_012536,NC_012537,NC_012538,NC_012539,NC_012 |

| |
|---|
| 754,NC_012755,NC_013225,NC_013226,NC_013227,NC_013228,NC_013229,NC_013230,NC_013396,NC_013397,NC_013398,NC_014236,NC_014237,NC_014238,NC_014239,NC_014240,NC_014241,NC_014511,NC_014512,NC_014513,NC_014514,NC_014522,NC_014523,NC_014598,NC_014599,NC_014600,NC_014601,NC_014602,NC_014708,NC_014709,NC_014710,NC_014714,NC_014715,NC_014716,NC_014717,NC_015126,NC_015127,NC_015128,NC_015129,NC_015130,NC_015877,NC_015878,NC_015879,NC_015880,NC_015881,NC_016874,NC_016875,NC_016876,NC_016879,NC_016880,NC_016881,NC_020439,NC_020440,NC_020441,NC_020442,NC_020447,NC_021541,NC_021543,NC_021545,NC_021551,NC_021580,NC_021581,NC_021589,NC_021590,NC_021625,NC_021626,NC_021630,NC_021631,NC_022553,NC_022554,NC_022555,NC_022620,NC_022626,NC_022627,NC_022633,NC_022634,NC_022639,NC_023420,NC_023486,NC_023487,NC_023488,NC_023491,NC_023492,NC_023813,NC_023814,NC_023815,NC_023816,NC_023819,NC_023820,NC_024503,NC_024504,NC_024505,NC_024506,NC_024507,NC_024916,NC_024917,NC_024918,NC_024919,NC_025485,NC_025486,NC_025487,NC_025488,NC_025493,NC_025801,NC_025802,NC_025803,NC_025804,NC_025808,NC_025845,NC_025846,NC_025847,NC_025848,NC_025849,NC_025850,NC_025851,NC_026825,NC_026826,NC_026827,NC_026828,NC_027533,NC_027534,NC_027535,NC_027539,NC_027553,NC_027554,NC_027567,NC_027568,NC_027569,NC_027572,NC_027574,NC_027803,NC_027808,NC_027809,NC_027811,NC_027812,NC_027816,NC_028465,NC_029904,NC_029911,NC_029912,NC_029913,NC_029914,NC_029917,NC_029918,NC_030158,NC_030159,NC_030160,NC_030161,NC_030162,NC_030163,NC_030405,NC_030406,NC_030412,NC_030413,NC_030414,NC_030415,NC_033782,NC_033783,NC_033784,NC_034168,NC_034169,NC_034170,NC_034171,NC_034172,NC_035935,NC_035936,NC_036468,NC_036469,NC_036470,NC_036471,NC_036476,NC_036477,NC_037570,NC_037571,NC_037572,NC_037573,NC_037574,NC_037578,NC_037579,NC_037580,NC_037581,NC_037582,NC_037583,NC_038564,NC_038565,NC_038568,NC_038570,NC_038574,NC_038575,NC_038582,NC_038584,NC_038588,NC_038592,NC_038594,NC_038595,NC_038600,NC_038604,NC_038605,NC_038610,NC_038614,NC_038615,NC_038620,NC_038624,NC_038625,NC_038629,NC_038630,NC_038634,NC_038635,NC_038636,NC_038637,NC_038640,NC_038641,NC_038645,NC_038648,NC_038649,NC_038652,NC_038657,NC_038660,NC_038661,NC_038662,NC_038664,NC_038665,NC_038945,NC_038948,NC_040408,NC_040409,NC_040413,NC_040414,NC_040440,NC_040443,NC_040444,NC_040445,NC_040447,NC_040472,NC_040473,NC_040476,NC_040478,NC_040479,NC_040499,NC_040501,NC_040502,NC_040503,NC_040504,NC_040506,NC_043180,NC_043182,NC_043183,NC_043184,NC_043185,NC_043190,NC_043368,NC_043369,NC_043370 |
| **Rhabdoviridae** |
| KC519324,KC685626,KP688373,NC_000855,NC_000903,NC_001542,NC_001560,NC_001615,NC_001652,NC_002251,NC_002526,NC_002803,NC_003243,NC_003746,NC_005093,NC_005974,NC_005975,NC_006429,NC_006942,NC_007020,NC_007642,NC_008514,NC_009527,NC_009528,NC_009608,NC_009609,NC_011532,NC_011558,NC_011568,NC_011639,NC_013135,NC_013955,NC_016136,NC_017685,NC_017714,NC_018381,NC_018629,NC_020803,NC_020804,NC_020805,NC_020806,NC_020807,NC_020808,NC_020809,NC_020810,NC_022580,NC_022581,NC_022755,NC_024473,NC_025251,NC_025253,NC_025255,NC_025340,NC_025341,NC_025342,NC_025353,NC_025356,NC_025358,NC_025359,NC_025362,NC_025364,NC_025365,NC_025371,NC_025376,NC_025377,NC_025378,NC_025382,NC_025384,NC_025385,NC_025387,NC_025389,NC_025391,NC_025393,NC_025394,NC_025395,NC_025396,NC_025397,NC_025399,NC_025400,NC_025401,NC_025405,NC_025406,NC_025408,NC_028230,NC_028231,NC_028232,NC_028234,NC_028236,NC_028237,NC_028239,NC_028241,NC_028244,NC_028246,NC_028255,NC_028266,NC_028867,NC_030451,NC_031079,NC_031083,NC_031093,NC_031215,NC_031216,NC_031225,NC_031227,NC_031232,NC_031236,NC_031240,NC_031268,NC_031272,NC_031273,NC_031276,NC_031278,NC_031282,NC_031283,NC_031301,NC_031303,NC_031305,NC_031690,NC_031691,NC_031955,NC_031957,NC_031958,NC_031988,NC_033701,NC_033705,NC_034240,NC_034443,NC_034447,NC_034448,NC_034449,NC_034450,NC_034451,NC_034454,NC_034508,NC_034529,NC_034530,NC_034531,NC_034533,NC_034534,NC_034535,NC_034536,NC_034537,NC_034538,NC_034539,NC_034540,NC_034541,NC_034542,NC_034543,NC_034544,NC_034545,NC_034546,NC_034548,NC_034549,NC_034550,NC_034551,NC_036390,NC_038236,NC_038275,NC_038276,NC_038277,NC_038278,NC_038279,NC_038280,NC_038281,NC_038282,NC_038283,NC_038284,NC_038285,NC_038286,NC_038287,NC_038755,NC_038756,NC_039020,NC_039021,NC_039200,NC_039201,NC_039202,NC_039206,NC_040532,NC_040599,NC_040602,NC_040664,NC_040669,NC_040786,NC_043065,NC_043066,NC_043067,NC_043525,NC_043538,NC_043648,NC_043649,Z93414 |

| Secoviridae |
| --- |
| NC_001632,NC_003003,NC_003004,NC_003445,NC_003446,NC_003495,NC_003496,NC_003502,NC_003509,NC_003544,NC_003545,NC_003549,NC_003550,NC_003615,NC_003621,NC_003622,NC_003623,NC_003626,NC_003628,NC_003693,NC_003694,NC_003738,NC_003741,NC_003785,NC_003786,NC_003787,NC_003788,NC_003791,NC_003792,NC_003799,NC_003800,NC_003839,NC_003840,NC_004439,NC_004440,NC_005096,NC_005097,NC_005266,NC_005267,NC_005289,NC_005290,NC_006056,NC_006057,NC_006271,NC_006272,NC_006964,NC_006965,NC_008182,NC_008183,NC_009013,NC_009032,NC_010709,NC_010710,NC_010987,NC_010988,NC_011189,NC_011190,NC_013075,NC_013076,NC_013218,NC_013219,NC_015414,NC_015415,NC_015492,NC_015493,NC_016443,NC_016444,NC_017938,NC_017939,NC_018383,NC_018384,NC_020897,NC_020898,NC_022004,NC_022006,NC_022798,NC_022799,NC_023016,NC_023017,NC_025479,NC_025480,NC_027915,NC_027926,NC_027927,NC_028139,NC_028146,NC_029036,NC_029038,NC_031763,NC_031766,NC_032270,NC_032271,NC_033492,NC_033493,NC_034214,NC_034215,NC_035214,NC_035215,NC_035218,NC_035219,NC_035220,NC_035221,NC_038320,NC_038744,NC_038759,NC_038760,NC_038761,NC_038762,NC_038763,NC_038764,NC_038765,NC_038766,NC_038767,NC_038768,NC_038862,NC_038863,NC_039072,NC_039073,NC_039077,NC_039078,NC_040399,NC_040400,NC_040416,NC_040417,NC_040586,NC_043076,NC_043385,NC_043388,NC_043411,NC_043447,NC_043448,NC_043684,NC_043685 |
| **Test-3a; Source: Virus-Host-DB; NCBI; GISAID** |
| **Alphacoronavirus** |
| AY518894,FJ938054,FJ938055,FJ938056,FJ938057,FJ938058,FJ938059,FJ938060,FJ938061,FJ938062,GU553361,D13096,GU553362,HM245926,HQ392469,HQ392470,HQ392471,HQ392472,JN183882,JN183883,JQ410000,JQ989272,DQ811787,LM645057,NC_002306,NC_002645,NC_003436,NC_005831,NC_009657,NC_009988,NC_010437,NC_010438,NC_018871,DQ848678,NC_022103,NC_023760,NC_028752,NC_028806,NC_028811,NC_028814,NC_028824,NC_028833,NC_030292,NC_032107,EU420137,NC_032730,NC_034972,NC_035191,NC_038861,EU420138,FJ938051,FJ938052,FJ938053 |
| **Betacoronavirus** |
| EU371561,EU371562,EU371563,EU371564,FJ415324,FJ425184,FJ425185,FJ425186,FJ425187,FJ425188,FJ425189,FJ647218,FJ647219,FJ647220,FJ647221,FJ647222,FJ647223,FJ647224,FJ647225,FJ647226,FJ647227,FJ882935,FJ882942,FJ882945,FJ882954,FJ882963,FJ884686,FJ938063,FJ938064,FJ938065,FJ938066,FJ938067,GQ153539,GQ153540,GQ153541,GQ153542,GQ153543,GQ153544,GQ153545,GQ153546,GQ153547,GQ153548,HM211098,HM211099,HM211100,HM211101,JF792616,JQ173883,JX169867,JX860640,JX869059,JX993987,JX993988,KC667074,KC776174,KC881005,KC881006,KF367457,KF569996,KF906249,KJ473821,NC_001846,NC_003045,NC_004718,NC_006213,NC_006577,NC_009019,NC_009020,NC_009021,NC_012936,NC_017083,NC_019843,NC_025217,NC_026011,NC_030886,NC_038294,NC_039207,AC_000192,AF201929,AY278488,AY278491,AY278554,AY278741,AY350750,AY357075,AY357076,AY394850,AY515512,AY864805,AY864806,DQ011855,DQ412042,DQ412043,DQ640652,DQ648794,DQ648856,DQ648857,DQ915164,EF065505,EF065506,EF065507,EF065508,EF065509,EF065510,EF065511,EF065512,EF065513,EF065514,EF065515,EF065516,EF424615,EF424616,EF424617,EF424618,EF424619,EF424620,EF424621,EF424622,EF424623,EF424624,EF446615,EU371559,EU371560,MG772933.1,MG772934.1,EPI_ISL_402131 |
| **Deltacoronavirus** |
| FJ376620,KJ481931,KJ567050,KJ601777,KJ601778,KJ601779,KJ601780,KJ769231,KM820765,KP981395,NC_011547,NC_011549,NC_011550,NC_016991,NC_016992,NC_016993,NC_016994,NC_016995,NC_016996,NC_039208 |
| **Gammacoronavirus** |
| AY646283,FN430414,FN430415,JF705860,KF793824,LN610099,NC_001451,NC_010646,NC_010800 |
| **Test-3b; Source: Virus-Host-DB; GISAID** |
| **Alphacoronavirus** |
| JQ989272,JQ410000,DQ811787,FJ938058,NC_022103,NC_028752,EU420137,NC_038861,FJ938051,FJ938056,FJ938059,NC_034972,NC_028811,HQ392471,FJ938057,NC_028824,NC_028814,FJ938060,HM245926,NC_028833 |

| |
|---|
| **Betacoronavirus** |
| EF065513,FJ882942,FJ425185,HM211100,GQ153540,NC_006213,GQ153543,EF424624,FJ647220,FJ938065,EPI_ISL_402131,FJ938066,AY278554,DQ915164,DQ011855,FJ882945,FJ647225,FJ425184,FJ415324,FJ882935 |
| **Deltacoronavirus** |
| FJ376620,KJ481931,KJ567050,KJ601777,KJ601778,KJ601779,KJ601780,KJ769231,KM820765,KP981395,NC_011547,NC_011549,NC_011550,NC_016991,NC_016992,NC_016993,NC_016994,NC_016995,NC_016996,NC_039208 |
| **Test-4; Source: Virus-Host-DB; NCBI; GISAID** |
| **Embecovirus** |
| AC_000192,EF424620,EF424621,EF424622,EF424623,EF424624,EF446615,FJ415324,FJ425184,FJ425185,FJ425186,AF201929,FJ425187,FJ425188,FJ425189,FJ647218,FJ647219,FJ647220,FJ647221,FJ647222,FJ647223,FJ647224,DQ011855,FJ647225,FJ647226,FJ647227,FJ884686,FJ938063,FJ938064,FJ938065,FJ938066,FJ938067,JF792616,DQ915164,JQ173883,JX169867,JX860640,KF906249,NC_001846,NC_003045,NC_006213,NC_006577,NC_012936,NC_026011,EF424615,EF424616,EF424617,EF424618,EF424619 |
| **Merbecovirus** |
| DQ648794,EF065505,EF065506,EF065507,EF065508,EF065509,EF065510,EF065511,EF065512,JX869059,KC667074,KC776174,KJ473821,NC_009019,NC_009020,NC_019843,NC_038294,NC_039207 |
| **Nobecovirus** |
| EF065513,EF065514,EF065515,EF065516,HM211098,HM211099,HM211100,HM211101,NC_009021,NC_030886 |
| **Sarbecovirus** |
| MG772933.1,MG772934.1,EPI_ISL_402131,FJ882935,FJ882942,FJ882945,FJ882954,FJ882963,GQ153539,GQ153540,GQ153541,GQ153542,GQ153543,GQ153544,GQ153545,GQ153546,GQ153547,GQ153548,JX993987,JX993988,KC881005,KC881006,KF367457,KF569996,NC_004718,AY278488,AY278491,AY278554,AY278741,AY350750,AY357075,AY357076,AY394850,AY515512,AY864805,AY864806,DQ412042,DQ412043,DQ640652,DQ648856,DQ648857,EU371559,EU371560,EU371561,EU371562,EU371563,EU371564 |
| **Test-5; Source: Virus-Host-DB; NCBI; GISAID** |
| **Embecovirus; Merbecovirus; Nobecovirus; Sarbecovirus** |
| same as Test-4 |
| **2019-nCoV** |
| EPI_ISL_402119,EPI_ISL_402130,EPI_ISL_402132,EPI_ISL_403928,EPI_ISL_403929,EPI_ISL_403930,EPI_ISL_403931,EPI_ISL_403932,EPI_ISL_403933,EPI_ISL_403934,EPI_ISL_403935,EPI_ISL_402120,EPI_ISL_403936,EPI_ISL_403937,EPI_ISL_403962,EPI_ISL_403963,EPI_ISL_404227,EPI_ISL_404228,MN908947.3,EPI_ISL_402121,EPI_ISL_402123,EPI_ISL_402124,EPI_ISL_402125,EPI_ISL_402127,EPI_ISL_402128,EPI_ISL_402129,EPI_ISL_404253,EPI_ISL_404895,EPI_ISL_405839 |
| **Test-6; Source: Virus-Host-DB; NCBI; GISAID** |
| **Sarbecovirus; 2019-nCoV** |
| same as Test-5 |

**Supplementary Table S3:** Accession IDs of sequences used in Test-1 to Test-6.
